# The splicing regulator Prp31 prevents retinal degeneration in *Drosophila* by regulating Rhodopsin levels

**DOI:** 10.1101/386433

**Authors:** Malte Lehmann, Sarita Hebbar, Behrens Sarah, Weihua Leng, Michaela Yuan, Sylke Winkler, Elisabeth Knust

## Abstract

Retinitis pigmentosa (RP) is a clinically heterogeneous disease affecting 1.6 million people worldwide. The second-largest group of genes causing autosomal dominant RP in human encodes regulators of the splicing machinery, but the molecular consequences that link defects in splicing factor genes to the aetiology of the disease remain to be elucidated. Mutations in PRPF31, one of the splicing factors, are linked to RP11. To get insight into the mechanisms by which mutations in this gene lead to retinal degeneration, we induced mutations in the *Drosophila* orthologue *Prp31*. Flies heterozygous mutant for *Prp31* are viable and develop normal eyes and retina. However, photoreceptors degenerate under light stress, thus resembling the human disease phenotype. *Prp31* mutant flies show a high degree of phenotypic variability, similar as reported for human RP11 patients. Degeneration is associated with increased accumulation of rhodopsin 1, both in the rhabdomere and in the cell body. In fact, reducing rhodopsin levels by raising animals in a carotenoid-free medium not only suppressed rhodopsin accumulation, but also retinal degeneration. In addition, our results underscore the relevance of eye color mutations on phenotypic traits, in particular whilst studying a complex process such as retinal degeneration.

**Article Summary:** Retinitis pigmentosa (RP) is a human disease affecting 1.6 million people worldwide. So far >50 genes have been identified that are causally related to RP. Mutations in the splicing factor PRPF31 are linked to RP11. We induced mutations in the *Drosophila* orthologue *Prp31* and show that flies heterozygous for *Prp31* undergo light-dependent retinal degeneration. Degeneration is associated with increased accumulation of the light-sensitive molecule, rhodopsin 1. In fact, reducing rhodopsin levels by dietary intervention suppressed retinal degeneration. We believe that this model will help to better understand the aetiology of the human disease.

## Introduction

Retinitis pigmentosa (RP; OMIM 268000) is a clinically heterogeneous set of retinal dystrophies, which affects about 1.6 million people worldwide. It often starts with night blindness in early childhood due to the degeneration of rod photoreceptor cells (PRCs), continues with the loss of the peripheral visual field caused by degeneration of rods (tunnel vision), and progresses to complete blindness in later life. RP is a genetically heterogeneous disease and can be inherited as autosomal dominant (adRP), autosomal recessive (arRP) or X-linked (xlRP) disease. So far >50 genes have been identified that are causally related to non-syndromic RP (see RetNet: http://www.sph.uth.tmc.edu/RetNet/disease.htm). Affected genes are functionally diverse. Some of them are expressed specifically in PRCs and encode, among others, transcription factors (e. g. *CRX*, an *otx*-like photoreceptor homeobox gene), components of the light-induced signalling cascade, including the visual pigment rhodopsin (Rho/*RHO* in *Drosophila*/human), or genes controlling vitamin A metabolism (e.g. *RLBP-1*, encoding Retinaldehyde-binding protein). Other genes are associated with the control of cellular homeostasis, for example *CRB1*, a gene required for the maintenance of polarity (Daiger *et al.* 2013; Daiger *et al.* 2014). Interestingly, the second-largest group of genes causing adRP, comprising 7 of 23 genes known, encodes regulators of the splicing machinery. So far, mutations in five PRPF (pre-mRNA processing factor) genes, PRPF31, PRPF4, PRPF6, PRPF8 and PRPF31, have been linked to adRP, namely RP18, RP70, RP60, RP13 and RP11, respectively. PAP1 (Pim1-associated protein) and SNRNP200 (small nuclear ribonuclearprotein-200), two genes also involved in splicing, have been suggested to be associated with RP9 and RP33, respectively (MAITA *et al.* 2004; Zhao *et al.* 2009) [reviewed in (Mordes *et al.* 2006; Poulos *et al.* 2011; Liu and Zack 2013; Ruzickova and Stanek 2016)]. The five *PRPF* genes encode components regulating the assembly of the U4/U6.U5 tri-snRNP, a major module of the pre-mRNA spliceosome machinery (Will and Luhrmann 2011). Several hypotheses have been put forward to explain why mutations in ubiquitously expressed components of the general splicing machinery show a dominant phenotype only in PRCs. One hypothesis suggests that PRCs with only half the copy number of a gene encoding a general splicing component cannot cope with the elevated demand of RNA-/protein synthesis required to maintain the exceptionally high metabolic rate of PRCs in comparison to other tissues. Hence, halving their gene dose eventually results in apoptosis. Although this model is currently favoured, other mechanisms, such as impaired splicing of PRC-specific mRNAs or toxic effects caused by accumulation of mutant proteins have been discussed and may contribute to the disease phenotype [discussed in (Mordes *et al.* 2006; Tanackovic *et al.* 2011; Scotti and Swanson 2016)].

The observation that all adRP-associated genes involved in splicing are highly conserved from yeast to human allows to use model organisms to unravel the genetic and cell biological functions of these genes in order to obtain mechanistic insight into the origin of the diseases. In the case of RP11, the disease caused by mutations in *PRPF31*, three mouse models have been generated by knock-in and knock-out approaches. Unexpectedly, mice heterozygous mutant for a null allele or a point mutation that recapitulates a mutation in the corresponding human gene did not show any sign of retinal degeneration in 12 and 18-month-old mice, respectively (Bujakowska *et al.* 2009). Further analyses revealed that the retinal pigment epithelium, rather than the PRCs, is the primary tissue affected in *Prpf31* heterozygous mice (Graziotto *et al.* 2011; Farkas *et al.* 2014). Morpholino-induced knock-down of zebrafish *Prpf31* results in strong defects in PRC morphogenesis and survival (Linder *et al.* 2011). Defects induced by retina-specific expression of zebrafish *Prpf31* constructs that encode proteins with the same mutations as those mapped in RP11 patients (called AD5 and SP117, respectively) were explained to occur by either haplo-insufficiency or by a dominant-negative effect of the mutant protein (Yin *et al.* 2011). In *Drosophila*, no mutations in the orthologue *Prp31* have been identified so far. RNAi-mediated knock-down of *Prp31* in the *Drosophila* eye results in abnormal eye development, ranging from smaller eyes to complete absence of the eye, including loss of PRCs and pigment cells (Ray *et al.* 2010).

We aimed to establish a meaningful *Drosophila* model for RP11-associated retinal degeneration in order to get better insights into the mechanisms by which *Prp31* prevents retinal degeneration. Therefore, we isolated two mutant alleles of *Prp31*, *Prp31*^*P17*^ and *Prp31*^*P18*^, which carry missense mutations affecting conserved amino acids. Flies heterozygous for either of these mutations are viable and develop normally. Strikingly, when exposed to constant light, these mutant flies undergo retinal degeneration, thus mimicking the disease phenotype of RP11 patients. Degeneration of mutant PRCs is associated with accumulation and abnormal distribution of the visual pigment rhodopsin, Rh1, in PRCs. Reduction of dietary vitamin A, a precursor of the chromophore 11-cis-3-hydroxyretinal, which is bound to opsin to generate the functional visual pigment rhodopsin, prevents accumulation of rhodopsin and retinal degeneration. From this we conclude that Rh1 accumulation/misdistribution is a major cause of retinal degeneration in *Prp31* heterozygous flies. We provide additional evidence for the strong influence of the genetic background on the expressivity of the mutant phenotype, a feature that has also been described for the human disease.

## Materials and Methods

### Fly strains and genetics

All phenotypic analyses were performed in age-matched males unless otherwise specified. Genotypes and genders are summarized in Supplemental Table 1. Flies were maintained at 25°C on standard yeast-cornmeal-agar food. To rule out differences in light sensitivity in the light-degeneration paradigm, we utilized white-eyed flies, bearing mutations in the *white* gene, either as general controls or in the mutant background. We tested two *white* alleles (*w\** and *w^1118^*). Molecular testing of these two alleles by PCR revealed that both *w\** and *w^1118^* carry a deletion of the transcriptional and translation start site of the *white* gene (Fig. S1A). However, whilst both lines respond to constant light exposure, *w^1118^* exhibits a drastic loss of photoreceptor cells, in that 75% of all ommatidia are damaged in *w^1118^* eyes following constant light exposure (Fig. S1B-E). In contrast, *w\** only exhibits modest changes in morphology, consistent with expected effects of constant light exposure. It has been recently reported that *w^1118^* is the most severely affected *w* allele in degeneration paradigms as compared to other alleles of *white* (Ferreiro *et al.* 2018). Furthermore, different strains of *w*^*1118*^ have been reported to exhibit varying phenotypes in terms of adult behaviour (Sun *et al.* 2009). Based on these data, we utilized *w\** as our general control, given its stereotypic response to constant light. The RNAi line (ID: 35131) for the *Prp31* gene was obtained from the Vienna *Drosophila* Resource Centre (VDRC, www.vdrc.at) (Dietzl *et al.* 2007). RNAi was induced either using *Rh1-Gal4* in combination with Dicer-2 expression, or with a transgene (*GMR-w^IR^)* (Lee and Carthew 2003) to assay degeneration in a non-pigmented background. *Df(3L)Exel6262* with deleted segment 71B3;71C1 (Parks *et al.* 2004)*, Df(3L)ED217* with deleted segment 70F4;71E1 and *Df(3L)ED218* with deleted segment 71B1 - 71E1 (Ryder *et al.* 2007) were obtained from the Bloomington Stock Centre. Since the deficiency lines carry a mini-*white* transgene due the way they were generated (Ryder *et al.* 2007), *cn bw* was recombined into these lines and all phenotypes were compared with *cn bw*. *gstD-GFP* (Sykiotis and Bohmann 2008) (gift from D. Bohman), recombined into *Prp31^18^,* deficiency lines or genetic controls were used as an indicator of oxidative stress signalling.

### Isolation of *Prp31* alleles by TILLING

To isolate point mutations in the *Prp31* locus (FlyBase ID: FBgn0036487) a library, of 2.400 fly lines with isogenized third chromosomes, which potentially carry point mutations caused by EMS treatment, was screened. Our approach targeted exon 1-3 of the *Prp31* locus containing two thirds (67%) of the coding sequence and including several predicted functional domains (the NOSIC (IPRO012976), the Nop (IPRO002687) and parts of the Prp31_C terminal (IPRO019175) domain), making use of two different PCR amplicons. A nested PCR approach was used, where the inner primers contain universal M13 tails that serve as primer binding sites of the Sanger sequencing reaction:

- amplicon1 (covers exon 1 and 2), outer primer, forward: TTCAATGAACCGCATGG, reverse: GTCGATCTTTGCCTTCTCC, inner / nested primer, forward: TGTAAAACGA CGGCCAGT-AGCAACGGTCACTTCAATTC, reverse: AGGAAACAGCTATGACCAT-GAAAGGGAATGGGATTCAG);
- amplicon 2 (covers exon 3), outer primer, forward: ATCGTGGGTGAAATCGAG, reverse: TGGTCTTCTCATCCACCTG, inner / nested primer, forward: TGTAAAACGA CGGCCAGT-AAGCTGCAGGCTATTCTCAC, reverse: AGGAAACAGCTATGACCAT-TAGGCATCCTCTTCGATCTG.

PCR-reactions were performed in 10 µl volume and with an annealing temperature of 57 °C, in 384 well format, making use of automated liquid handling tools. PCR fragments were sequenced by Sanger sequencing optimized for amplicon re-sequencing in a large-scale format (Winkler *et al.* 2005; Winkler *et al.* 2011). Primary hits, resembling sequence variants, which upon translation result in potential nonsense and missense mutations or affect a predicted splice site, were verified in independent PCR amplification and Sanger sequencing reactions. Two of the four lines, named *Prp31*^*P17*^ and *Prp31*^*P18*^, were recovered from the living fly library and crossed for three generations to control, white-eyed (*w^*^*) flies to reduce the number of accompanying sequence variations. The removal of the markers of the original, mutagenized chromosome (*ru st e ca*) by the above outcrossing was verified as follows: the isolated alleles (males) were crossed to the original line (*ru st e ca*) and checked for the phenotypes associated with homozygous conditions of *roughoid* (*ru*; eye appearance)*, scarlet* (*st*; eye colour), *ebony* (*e*; body colour), *claret* (*ca*; eye color).

### Experimental light conditions

Flies were reared in regular light conditions defined as 12 hours of light (approx. 900-1300 lux)/12 hours of darkness. For the light-induced degeneration setting, flies (2 days of age) were placed at 25°C for 7 days in an incubator dedicated for continuous, high intensity light exposure (Johnson *et al.* 2002). High intensity light was defined by intensity of 1200-1300 lux measured using an Extech 403125 Light ProbeMeter (Extech Insturments, USA) with the detector placed immediately adjacent to the vial and facing the nearest light source. To reduce blue-green light in this setting, a customized box bounded by filters, which block blue-green light (shown in Fig. 3A) and face the light source in the incubator, was used. Light intensity was determined by measuring light counts using a USB spectrometer (Ocean Optics, USA). At the end of 7 days, fly heads were processed for sectioning. For immunostaining and western blotting, flies (1 day) reared under regular light were processed as described below.

### Vitamin A depletion

For vitamin A depletion experiments, animals were raised and maintained from embryonic stages onward on carotenoid free food (10% dry yeast, 10% sucrose, 0.02% cholesterol, and 2% agar) as described (Pocha *et al.* 2011).

### Transmission electron microscopy

Fixation of adult eyes, semi-thin sections and ultra-thin sections for transmission electron microscopy was performed as described (Mishra and Knust 2013). 1-2 µm semi-thin sections were stained with toluidine blue (1% / sodium tetraborate). 70nm ultrathin sections were imaged using a Morgagni 268 TEM (100kV) electron microscope (FEI Company), and images were taken using a Side-entry Morada CCD Camera (11 Megapixels, Olympus).

### Quantification of Degeneration

Toluidine blue stained semi-thin sections were imaged with a 63x Plan Apo oil objective (N.A. =1.4) on AxioImager.Z1 (Zeiss, Germany), fitted with AxioCamMRm camera, and analysed using the AxioVision software (Release 4.7). Quantification of degeneration was performed as described (Bulgakova *et al.* 2010). Briefly, the number of detectable rhabdomeres in each ommatidium was recorded from approximately 60-80 ommatidia per section and at least three eyes from different individuals were analysed. In case of degeneration, fewer ommatidia were counted since most of the tissue had degenerated.

### Immunostaining of adult retina and confocal imaging

Adult eyes were dissected and fixed in 4% formaldehyde. They were then processed either directly for immunostaining of the whole eye after removal of the lens, or for cryosectioning. For sectioning, sucrose treatment and embedding of the tissues in Richard-Allan Scientific NEG-50^TM^ (Thermo Fisher Scientific, UK) tissue embedding medium was done (Mishra and knust 2013). Eyes were cryosectioned at 12µm thickness at -21°C. Sections were air-dried and then subjected to immunostaining, which was done as described previously (Spannl *et al.* 2017). Antibodies used were rabbit anti-GFP (1:500; A-11122; Thermo Fisher Scientific, UK), mouse anti-Rh 1 (1:50; 4C5) and mouse anti-Na^+^-K^+^-ATPase (1:100; a5), both from Developmental Studies Hybridoma Bank (DSHB), University of Iowa, USA. 4C5 [http://dshb.biology.uiowa.edu/4C5] and a5 [http://dshb.biology.uiowa.edu/a5] were deposited to the DSHB by de Coet, H.G./Tanimura, T., and by Fambrough, D.M., respectively. Alexa-Flour conjugated secondary antibodies (1:200, Thermo Fisher Scientific, UK) were used. DAPI (4’,6-Diamidino-2-Phenylindole, Dihydrochloride; Thermo Fisher Scientific, UK) was used to label nuclei in tissue sections and Alexa-Fluor-555–phalloidin (Thermo Fisher Scientific, UK) was used to visualise F-actin enriched rhabdomeres. Sections and whole mounts were imaged with an Olympus Fluoview 1000 confocal microscope using an Olympus UPlanSApochromat 60x Oil objective (N.A. =1.35). They were subsequently visualized in Fiji (Schindelin *et al.* 2012) and corrected for brightness and contrast only.

### Western blotting

Five fly heads from each genotype were homogenized in 10 µL of 4x SDS-PAGE sample buffer (200 mM Tris-HCl pH 6.8, 20% Glycerol, 8% SDS, 0.04% Bromophenol blue, 400 mM DTT). After dilution with RIPA buffer (150 mM sodium chloride, 1% Triton X-100, 0.5% sodium deoxycholate, 0.1% SDS, 50 mM Tris pH 8), lysates were heated at 37°C for 30 min. Lysates equivalent to 2.5 heads were loaded and run on a 15% acrylamide gel, and the proteins transferred onto a membrane (Nitrocellulose Blotting Membrane 10600002; GE Healthcare Life Sciences; PA; US). Primary antibodies were incubated overnight at 4°C and included anti-Rh1 (4C5; 1:500) and anti-ß-Tubulin (E7; 1:5,000), both from Developmental Studies Hybridoma Bank (DSHB), University of Iowa, USA. 4C5 [http://dshb.biology.uiowa.edu/4C5] and E7 [http://dshb.biology.uiowa.edu/tubulin-beta-_2] were deposited to the DSHB by de Coet, H.G./Tanimura, T., and by M. McCutcheon/ S. Carroll respectively. As secondary antibody IRDye 800CW goat anti-Mouse IgG (1:15,000; LI-COR Biotechnology; NE; US) was used for an 1 h incubation. The fluorescent signal from the dry membrane was measured using a LI-COR Odyssey Sa Infrared Imaging System 9260-11P (LI-COR Biotechnology). The intensity of the bands was analysed using the Image Studio Ver 4.0 software. The reported value in Fig. 7 is obtained following normalization of the intensity values for Rh1 with the corresponding Tubulin intensity values and the number of heads loaded onto the gel.

### Figure panel preparation

All figure panels were assembled using Adobe Photoshop CS5.1 and Adobe Illustrator CS3 (Adobe Systems, USA). Statistical analyses and graphs were generated using GraphPad Prism (GraphPad Software, Inc, USA) and Microsoft Excel. For protein sequence visualization, Illustrator of Biological Sequences (IBS; (Liu *et al.* 2015)) software package was used.

## Results

### Two *Prp31* alleles discovered by TILLING

It was recently shown that RNAi-mediated knockdown of *Drosophila Prp31* in the eye using eye-specific Gal4-lines (*eyeless* (*ey*)-Gal4 or GMR-Gal4) results in abnormal eye development, ranging from smaller eyes to complete absence of the eye, including loss of photoreceptor cells (PRCs) and pigment cells (Ray *et al.* 2010). Since both Gal4 lines are expressed throughout eye development, some of the defects observed could be the result of impaired development, for example as a consequence of defective cell fate specification or eye morphogenesis.

We aimed to establish a more meaningful *Drosophila* model for RP11-associated retinal degeneration, a human disease associated with mutations in the human orthologue *Prpf31*, which would allow a deeper insight into the role of this splicing factor in the origin and progression of the disease. Therefore, we set out to isolate specific mutations in *Drosophila Prp31* by TILLING (**T**argeting **I**nduced **L**ocal **L**esions **IN G**enomes), following a protocol described recently (Spannl *et al.* 2017). In total, 2.400 genomes of EMS (ethyl methanesulfonate)-mutagenized flies were screened for sequence variants in two different amplicons of *Prp31*. Four sequence variants were identified, which were predicted to result in potentially deleterious missense mutations. Two of the four lines, named *Prp31*^*P17*^ and *Prp31*^*P18*^, were recovered from the living fly library and crossed for three generations to control, white-eyed (*w^*^*) flies to reduce the number of accompanying sequence variations. We outcrossed the mutants with white-eyes flies rather than with wild-type, red-eyed flies to generate a sensitised background for light-dependent degeneration experiments, since presence of the pigment granules surrounding each ommatidium contributes towards lower sensitivity to light (Stark and Carlson 1984). *Prp31*^*P18*^ was viable as homozygotes and in trans over any of three deficiencies, which remove, among others, the *Prp31* locus (Fig. 1A). In contrast, no homozygous *Prp31*^*P17*^ flies were obtained. However, *Prp31*^*P17*^ was viable in trans over *Prp31*^*P18*^ and over *Df(3L)ED217*. This suggests that the lethality was due to a second site mutation, which was not removed despite extensive out-crossing. We noticed that out-crossing *Prp31*^*P17*^ and *Prp31*^*P18*^ did not remove *scarlet* (*st*), one of the markers of the original, mutagenized chromosome (*ru st e ca*). Therefore, the correct genotypes of the two mutant lines are *w\**; *Prp31*^*P17*^, *st^1^* and *w\**; *Prp31*^*P18*^, *st*^*1*^. For simplicity, we will refer to them as *Prp31*^*P17*^ and *Prp31*^*P18*^ throughout the text.

**Figure 1:**
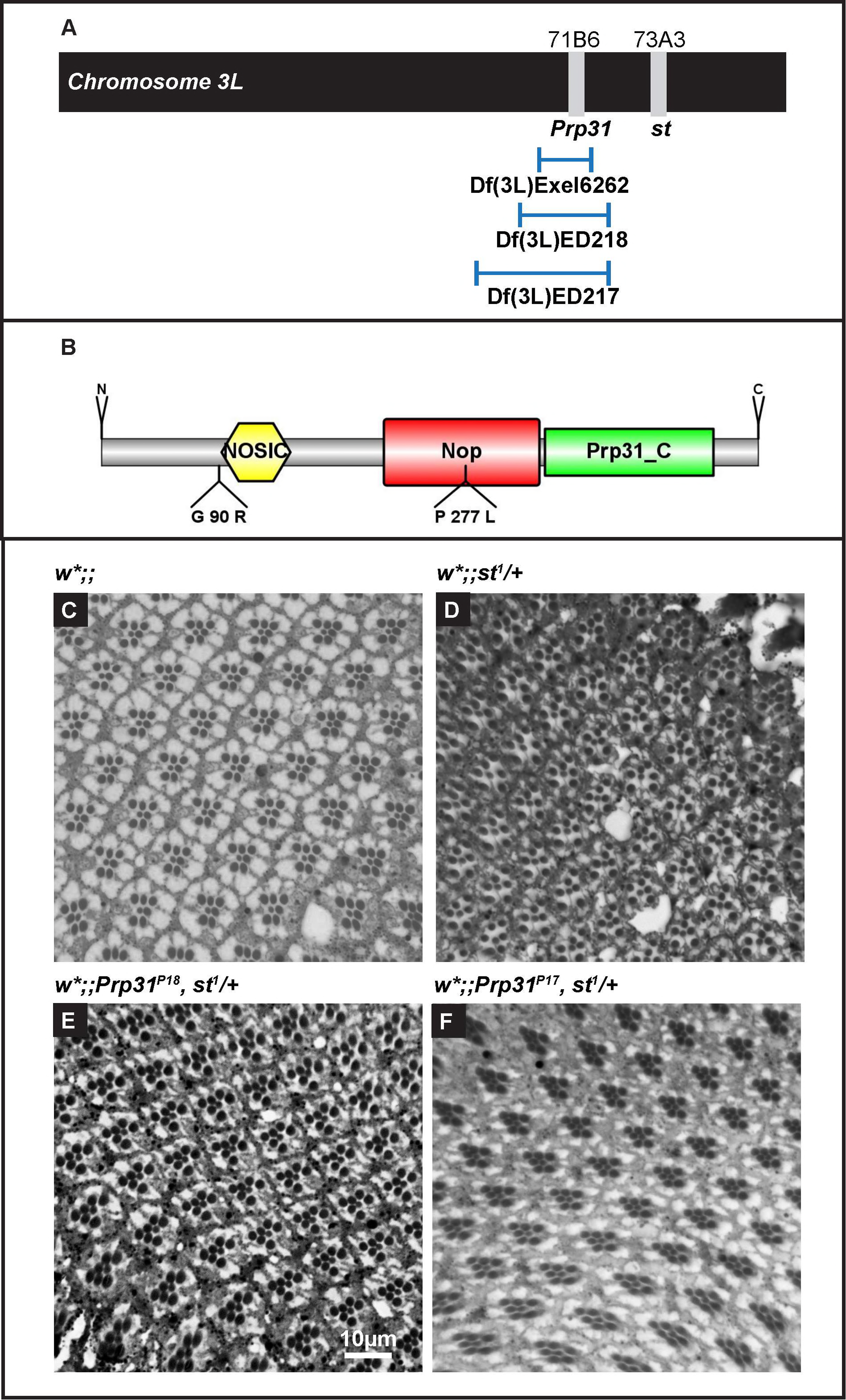
*Prp31* mutant flies have no gross morphological abnormalities at eclosion. (A). Schematic of chromosome arm 3L. *Prp31* and *scarlet* (*st*) are situated 2 cM apart (3-42 and 3-44, respectively; cytological positions 71B6 and 73A3, respectively; www.flybase.org). In both *Prp31* mutant alleles the marker *st^1^* from the original mutagenized chromosome (*ru st e ca*) is retained. The three deficiencies used cover the *Prp31* locus, but not the *st* locus. (B). Schematic overview of the Prpf31 protein. The figure is drawn to scale using IBS (Liu *et al.* 2015). Domains described here are indicated. The two *Prp31* alleles used here carry non-conservative missense mutations, G90R in *Prp31*^*P17*^ and P277L in *Prp31*^*P18*^. (**C-F**) Representative bright-field images of Toluidine-blue stained semi-thin sections of eyes of *w\** (C), *w*;; st^1^/+*(D), *w*;;Prp31^P18^, st^1^/+* (E), and *w*;;Prp31^P17^, st^1^/+* (F). Upon eclosion, flies were kept for two days under regular light conditions. Note that the number and stereotypic arrangement of photoreceptor cells within the mutant ommatidia are not affected. Scale bar = 10µm.

The molecular lesions in the two *Prp31* alleles were mapped in the protein coding region. *Drosophila* PRP31 is a protein of 501 amino acids, which contains a NOSIC domain (named after the central domain of Nop56/SIK1-like protein), a Nop (**N**ucle**o**lar **p**rotein) domain required for RNA binding, a PRP31_C-specific domain and a nuclear localization signal, NLS. *Prp31*^*P17*^ contained a point mutation that resulted in a non-conservative glutamine to arginine exchange (G90R) N-terminal to the NOSIC domain. *Prp31*^*P18*^ contained a non-conservative exchange of a proline to a leucine residue in the Nop domain (P277L) (Fig. 1B). Both mutations affect amino acids that are conserved between the fly and the human protein (Suppl. Fig. S2).

### Flies hetero- or hemizygous for *Prp31* undergo light-dependent retinal degeneration

Homo- and heterozygous *Prp31*^*P18*^ and heterozygous *Prp31*^*P17*^ animals raised and kept under regular light/dark cycles (12h light/12h dark) have eyes of normal size. Histological sections revealed normal numbers of PRCs per ommatidium (distinguished by the number of rhabdomeres) and a normal stereotypic arrangement of PRCs (Fig. 1C-F and Suppl. Fig. S2B). This indicates that the development of the retina was not affected by these mutations. However, PRCs of *Prp31*^*P17*^/+, *Prp31*^*P18*^/+ and *Prp31*^*P18*^/*Prp31*^*P18*^ flies showed clear signs of retinal degeneration when exposed to constant light for several days, manifested by a complete or partial loss of rhabdomeric integrity (Fig. 2A-D and Suppl. Fig. S2B’). We used the number of surviving rhabdomeres as an indicator of the severity of degeneration (Fig. 2E). When exposed for 7 days to constant light, *w\** mutant control flies exhibited some retinal degeneration, with 82% of all ommatidia still displaying the full complement of rhabdomeres. This phenotype is less severe than that reported for *w^1118^* flies (Chen *et al.* 2017; Ferreiro *et al.* 2018) (Fig. S1). *Prp31* mutant flies showed more severe PRC degeneration, with only about 48% of ommatidia having the full complement of seven PRCs (Fig. 2E). The degree of degeneration observed in *Prp31* alleles is less severe and more variable than that observed in the well-established RP12 disease model induced by mutations in the gene *crumbs* (*crb*) (Johnson *et al.* 2002; Chartier *et al.* 2012; Spannl *et al.* 2017). In the two *crb* alleles *crb^11A22^* and *crb^p13A9^* only 5 to 11% of all ommatidia displayed 7 rhabdomeres upon exposure to constant light, respectively (Fig. 2E).

**Figure 2:**
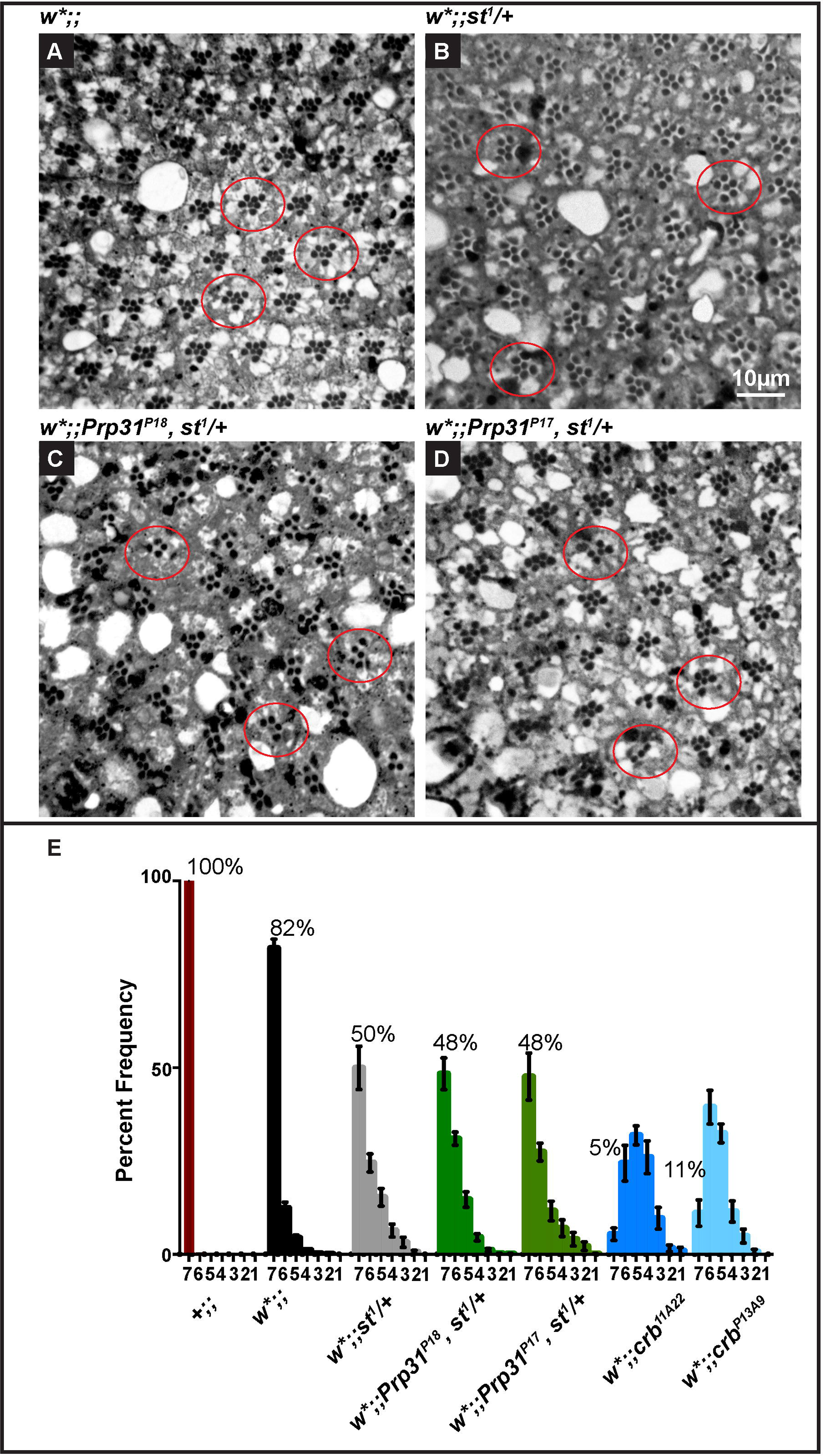
PRCs of heterozygous *Prp31*^*P17*^ or *Prp31*^*P18*^ flies undergo light-dependent degeneration. (**A-D**) Representative bright-field images of Toluidine-blue stained semi-thin sections of eyes of *w\** (A), *w*;; st^1^/+*(B), *w*;;Prp31^P18^, st^1^/+* (C), and *w*;;Prp31^P17^, st^1^/+* (D). Upon eclosion, flies were kept for two days under regular light conditions and then subjected to a degeneration paradigm of 7 days of continuous, high intensity light exposure. Whereas in w* (A) most ommatidia (red outline) display 7 rhabdomeres indicative of the 7 PRCs, *w*;; st^1^/+* and mutant ommatidia (B-D, red outlines) display fewer rhabdomeres per ommatidium indicative of degeneration. Scale bar = 10µm. (E) Quantification of retinal degeneration as indicated by the number of surviving rhabdomeres observed upon high intensity, continuous light exposure. Bars represent mean ± s.e.m. (a minimum of n=60 ommatidia from eyes of 3 biological replicates) of the percent frequency of ommatidia displaying 1-7 rhabdomeres (Y-axis). Genotypes are indicated below. Numbers on the graphs indicate the mean number of ommatidia displaying the full complement of 7 rhabdomeres.

Surprisingly, while about 18% of all ommatia in *w\** mutant control flies had less than seven rhabdomeres, this number was increased to 50% in the second genetic control, *w*;;st^1^/+* (Fig. 2E), suggesting that *st^1^* is a dominant enhancer of *w\**, at least with respect to retinal degeneration. This raised the question whether the degeneration observed in the two *Prp31* lines used is due to the mutation in *Prp31*, rather than to the mutations in *w* and *st*. To address this question, we reduced the intensity of blue/green light during light exposure (Fig. 3A), thereby minimising detrimental effects induced as a result of photolysis of rhodopsin, a known trigger of apoptosis (Stark and Carlson 1984; Stark *et al.* 1985). When exposed to filtered light with reduced blue/green intensity, neither *w\** nor *w*;; st^1^/+* ommatia displayed any sign of degeneration (Fig. 3B) and almost 100% of ommatidia showed the full complement of seven rhabdomeres. This is in stark contrast to results obtained under higher light intensity exposure, under which *w*;;* and *w*;;st^1^/+* displayed only 82% and 50% intact ommatidia, respectively (Fig. 3B). From this we concluded that the damage observed in eyes lacking pigments (*w*;;* and *w*;;st^1^/+*) is caused by high intensity light. This conclusion was corroborated by virtually no loss of rhabdomeres in wild-type (pigmented) eyes exposed to light (Suppl. Fig. S3). In contrast, in *Prp31*^*P18*^ heterozygous mutant flies exposed to lower blue/green light intensities still about 20% of all ommatidia displayed less than seven rhabdomeres, compared to 52% observed upon high intensity light exposure (Fig. 3B) Similarly, retinal degeneration is only slightly lowered in *crb* mutants at reduced blue green light, from 95% defective ommatidia to 80% (Fig. 3B). Another characteristic sign of light-induced tissue damage in white-eyed flies is the formation of holes or lacunae (Ferreiro *et al.* 2018). In fact, fewer holes were observed upon exposure to lower intensity light in the tissue (see Suppl. Fig. S3). Taken together, these results suggest i) that in flies lacking screening pigments high-intensity light induces tissue damage, i. e. PRC degeneration and formation of lacunae, which can be prevented by filtering-out high-energy wavelengths; and ii) that light-dependent retinal degeneration in *Prp31* mutants is due to the mutations in *Prp31*, and that the genetic background (*w*;;st^1^/+*) only marginally contributes to the degree of degeneration observed.

**Figure 3:**
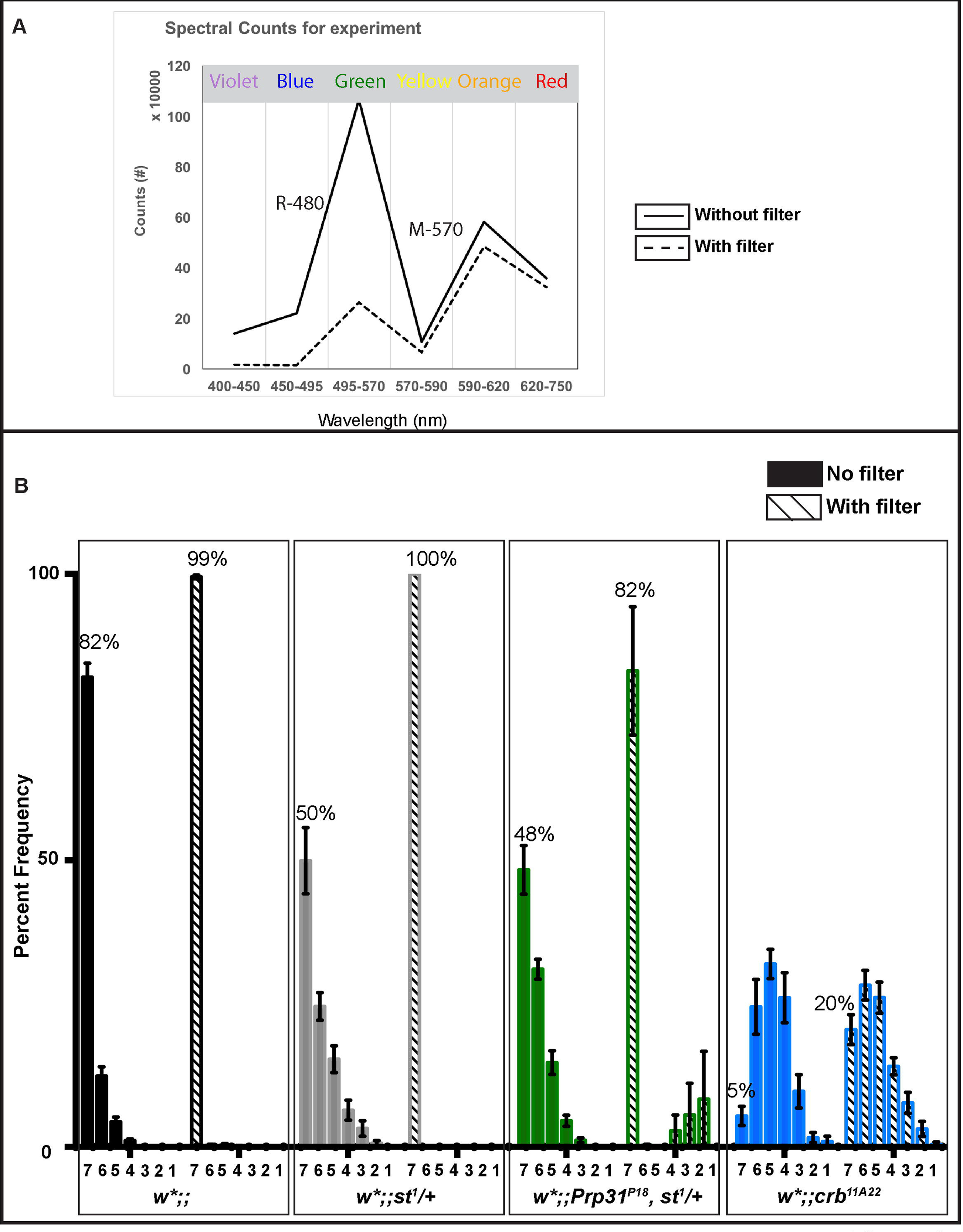
Reduced blue-green light intensitiy strongly reduces damage in *w^*^;;st/+* eyes. (A) Intensity profile (in counts as measured by a spectrometer) of light for the wavelength range (in nanometres) on the X-axis (the corresponding colour indicated above). The solid line represents the profile for the routine light degeneration paradigm, whereas the dashed line represents the intensity profile obtained when using a filter. Note that especially the intensity of the blue-green light is strongly reduced when using of a filter. (B) Quantification of retinal degeneration as indicated by the number of surviving rhabdomeres observed upon exposure to continuous, high-intensity light. Bars represent mean ± s.e.m. (a minimum of n=60 ommatidia from eyes of 3 biological replicates) of the percent frequency of ommatidia displaying 1-7 rhabdomeres (Y-axis). Genotypes are indicated below. For each genotype, the solid bar indicates surviving rhabdomeres under routine light-degeneration paradigm (corresponding to the solid line intensity profile in A), whereas the striped bar indicates surviving rhabdomeres upon reduced light intensity exposure (dashed line intensity profile in A).

To further confirm that the degeneration phenotype observed in *Prp31*^*P18*^ and *Prp31*^*P17*^ heterozygous flies is due to mutations in *Prp31* rather than to a mutation in *st*, we applied additional strategies to perturb *Prp31*. These included the use of *RNAi*-mediated knockdown of *Prp31* and of three deficiencies that remove *Prp31* but leave the *st* locus intact (Fig. 1A). First, we knocked down *Prp31* by overexpressing *Prp31* RNAi, mediated by *Rh1*-Gal4, which drives expression late in development, from 70% pupal development into adulthood (Kumar and Ready 1995). Thereby, we can rule out any effects on PRC specification or morphogenesis induced by loss of *Prp31*. To remove screening pigments, a second transgene was introduced into this background, called GMR-*w*^*IR*^, which expresses *white* RNAi under the control for the GMR-promoter. When exposed to light, the retina of corresponding control flies showed only minor morphological changes (Fig. 4A). However, the induction of *Prp31 RNAi* by *Rh1-Gal4* resulted in clear signs of degeneration upon light exposure, such as loss of rhabdomeres and accumulation of intensely stained structures reminiscent to apoptotic features (Fig. 4B). In fact, while 71% of control ommatidia revealed 7 identifiable rhabdomeres, this number decreased to 48% upon induction of *Prp31 RNAi* (Fig. 4C). As a second alternative strategy to study the role of *Prp31* in retinal degeneration we analysed the phenotype of three deficiency lines that cover the *Prp31* locus (see Fig. 1A). Since these deficiencies carry a *w*^+^-minigene, their retinal phenotype (and that of the respective control) was studied in a *w\**; *cn bw* mutant background in order to remove all screening pigments. *Df(3L)Exel6262/+*, *Df(3L)ED217/+*, and *Df(3L)ED218/+* flies exhibited the same degree of retinal degeneration as *Prp31*^*P17*^ or *Prp31*^*P18*^ heterozygous flies (Fig. 5), with only about 20% of their ommatidia showing seven rhabdomeres. Similar to the *Prp31* alleles, these deficiency lines had no obvious effects on retinal development (Suppl. Fig. S4A-D). Degeneration was also observed in hemizygous *Prp31* flies (*Prp31*^*P18*^/*Df (3L)217* and *Prp31*^*P17*^/*Df (3L)217*) (Suppl. Fig. S4E, F).

**Figure 4:**
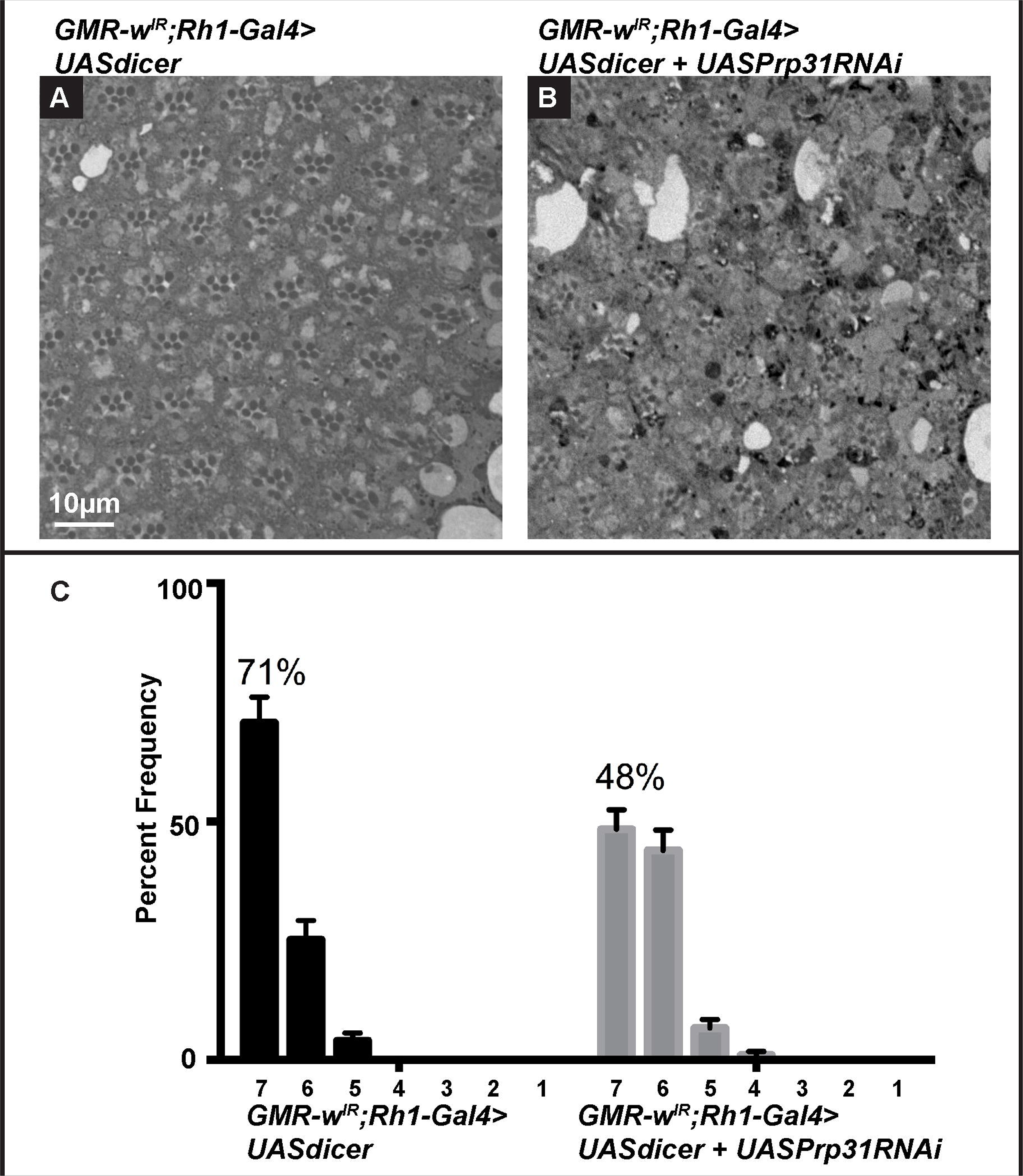
RNAi-mediated knock-down of *Prp31* results in light-dependent retinal degeneration. (**A-B**) Representative bright-field images of Toluidine-blue stained semi-thin sections of eyes of *GMR-w^IR^;Rh1-Gal4>UAS dicer* (A; control) and *GMR-w^IR^;Rh1-Gal4>UAS dicer + UAS Prp31RNAi* (**B**; *Prp31 RNAi*). Upon eclosion, flies were kept for two days under regular light conditions and then subjected to a degeneration paradigm of 7 days of continuous, high-intensity light exposure. In case of *Prp31 RNAi*, fewer ommatidia with 7 rhabdomeres are seen. Scale bar= 10µm (C) Quantification of retinal degeneration as indicated by the number of surviving rhabdomeres observed upon high intensity, continuous light exposure. Bars represent mean ± s.e.m. (a minimum of n=60 ommatidia from eyes of 3 biological replicates) of the percent frequency of ommatidia displaying 1-7 rhabdomeres (X-axis). Genotypes are indicated below. Whilst 71% of control ommatidia have 7 rhabdomeres, this number is reduced to 48% in the knockdown of *Prp31* by RNAi.

**Figure 5:**
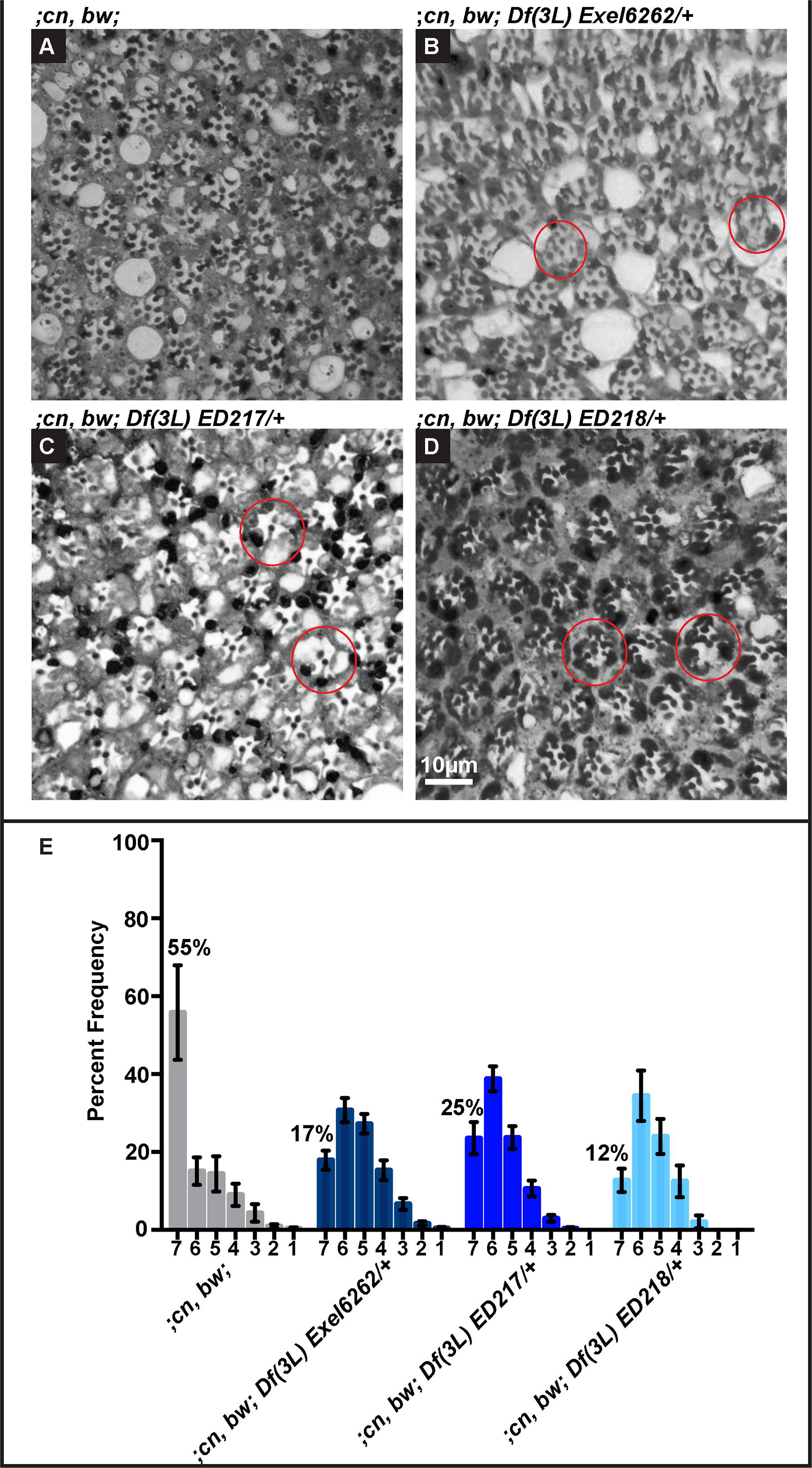
Flies heterozygous for deficiencies that cover *Prp31*, but not the *scarlet* locus, undergo light-dependent degeneration. (**A-D**) Representative bright-field images of Toluidine-blue stained semi-thin sections of eyes of males of *;cn, bw;* (A), *;cn, bw; Df (3L) Exel 6262/+ (B),; cn, bw; Df (3L) ED217/+* (C), and *;cn, bw; Df (3L) ED218/+* (D). Upon eclosion, flies were kept for two days under regular light conditions and then subjected to a degeneration paradigm of 7 days of continuous, high intensity light exposure. Scale bar= 10µm. (**E**) Quantification of retinal degeneration as indicated by the number of surviving rhabdomeres observed upon high intensity, continuous light exposure. Bars represent mean ± s.e.m. (a minimum of n=60 ommatidia from eyes of 3 biological replicates) of the percent frequency of ommatidia displaying 1-7 rhabdomeres (X-axis). Genotypes are indicated below.

Transmission electron microscopy (TEM) was used to further describe the ultrastructural features of degenerative phenotypes (Fig. 6). Hallmarks of degeneration include loss of rhabdomeral integrity, the complete loss of rhabdomeres in some PRCs, and the accumulation of electron dense aggregates. These features were mostly absent in eyes of *w\** flies and occur only to some extent in *w*;;st^1^/+* retina (Fig. 6A, B). In contrast, these attributes of degeneration were clearly identifiable and more pronounced in the retina of heterozygous and hemizygous *Prp31* flies (Fig. 6C-E). As mentioned above, degeneration in *crb* mutant eyes kept under the same conditions was more severe, as revealed from the complete loss of rhabdomeric integrity in all ommatidia and the accumulation of electron dense aggregates (Fig. 6F).

**Figure 6:**
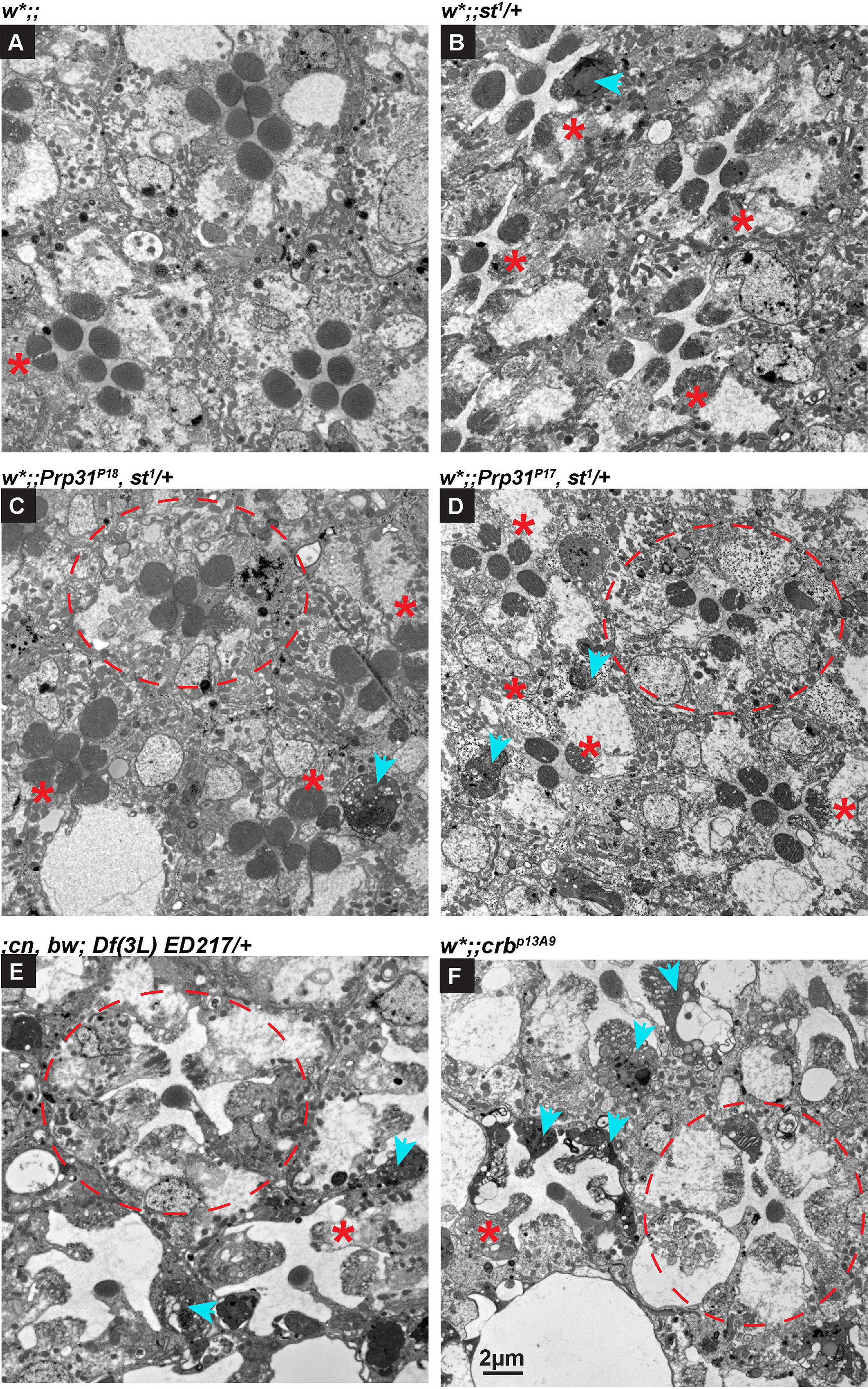
Hallmarks of degeneration in heterozygous *Prp31* mutants alleles revealed by TEM. (**A-F**) are representative transmission electron microscopy images of 70 nm sections of eyes of *w\** (A), *w*;; st^1^/+*(B), *w*;;Prp31^P18^, st^1^/+* (C), and *w*;;Prp31^P17^, st^1^/+* (D) *;cn, bw; Df (3L) ED217/+* (E), and *w*;; crb^p13A9^* (F) males. Upon eclosion, flies were kept for two days under regular light conditions and then subjected to a degeneration paradigm of 7 days of continuous, high intensity light exposure. Seven rhabdomeres are visible in the 3 ommatidia of the genetic controls (A-B). However, some rhabdomeres appear smaller or have lost their stereotypic appearance due to the loss of the microvillar structure (red asterisk). In heterozygous *Prp31* mutants (C-D) and in *Df (3L) ED217*/+ (E), some ommatidia (red outline) with PRCs lacking some rhabdomeres are obvious. All other ommatidia have PRCs with smaller or disintegrating rhabdomeres (red asterisk). The presence of large aggregates of electron dense material, another hallmark of degeneration, is more pronounced in perturbations of the *Prp31* gene (C-E, blue arrowheads). In *crb^13A9^* (F) all the above aspects of degeneration are visible, but. degeneration appears to be more severe than in the *Prp31* mutants. Scale bar = 2µm.

To summarise, data presented here support the conclusion that loss of one copy of the *Prp31* locus causes light-induced retinal degeneration.

### *Prp31* mutant photoreceptor cells show increased rhodopsin accumulation

A common cause of retinal degeneration, both in flies and in mammals, is abnormal localization/levels of the visual pigment rhodopsin1 (Rh1) (Hollingsworth and Gross 2012; Xiong and Bellen 2013). Therefore, we asked if the degeneration observed in *Prp31* mutant retinas is associated with altered Rh1 localization/levels. Rh1, encoded by *ninaE*, is the most abundant rhodopsin expressed in the outer PRCs R1-R6 (Ostroy *et al.* 1974; Harris *et al.* 1976). In control flies raised under regular light conditions (12h light/12h dark), Rh1 was concentrated in the rhabdomeres. As reported previously, Rh1 either fills the entire rhabdomere, forms a crescent-shaped pattern, or is restricted to the base or the lateral edges of the rhabdomere (Orem *et al.* 2006; Chinchore *et al.* 2009; Mitra *et al.* 2011; Xiong *et al.* 2012; Wang *et al.* 2014; Chen *et al.* 2017). Differences in localization have been attributed to inconsistency in antibody penetration due to the membrane-dense rhabdomeric structure (Xiong *et al.* 2012). The staining was more consistent when analysed in whole mount preparations. Here, Rh1 is more uniformly distributed, outlining the rhabdomeric structure along its length (Fig 7A’). Besides the rhabdomeric localization, Rh1 could be detected in cytoplasmic punctae (blue arrows in Fig. 7 and Suppl. Fig. S5,). This intracellular pool of Rh1 represents presumably internalized Rh1 following light exposure (Satoh and Ready 2005), since these flies were raised with 12 hours light and 12 hours darkness. PRCs of adult flies heterozygous for *Prp31* exhibited increased accumulation of Rh1 in the rhabdomeres in comparison to genetic controls (Fig. 7C, C’). Increased Rh1 immunostaining was observed in mutants independent of light conditions (Fig. S6).

**Figure 7:**
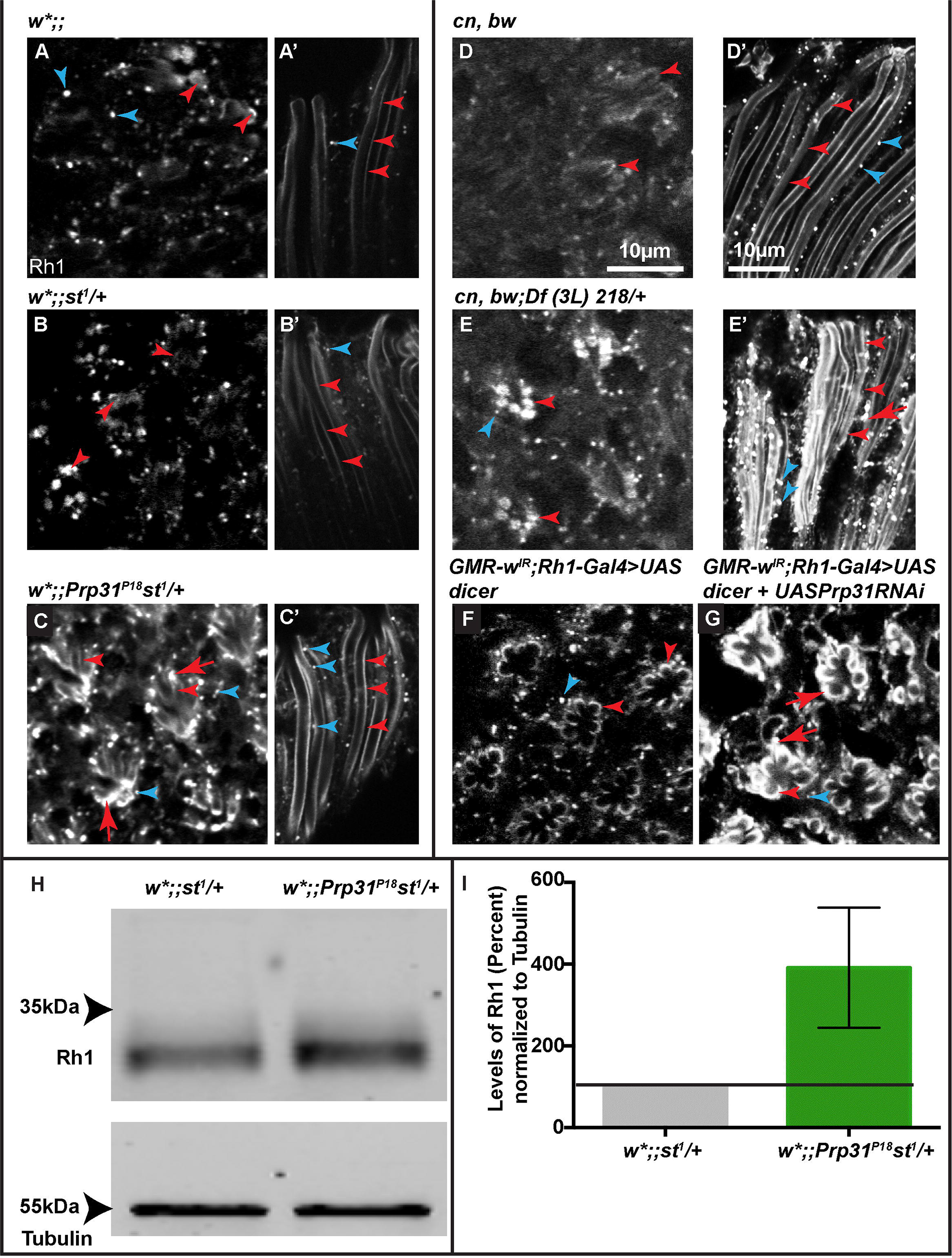
Increased Rhodopsin accumulation is associated upon perturbation of *Prp31*. Representative confocal images of 1µm optical sections from 12µm cross-sections (A-G), or whole mounts (A’- E’) of eyes of adult males with the genotypes indicated, stained with anti-Rh1. Red arrowheads indicate the rhabdomere, depicted in cross-sections (A-G) and along its length (A’-G’), with the distal end directed towards the top and the proximal end directed towards the bottom. Rh1 staining is more intense in the rhabdomeric membrane of *w*;;Prp31^P18^, st^1^/+* (C) as compared to controls, *w\** (A), *w*;; st^1^/+* (B). Increased intensity of Rh1 staining along the rhabdomeres and in sub-rhabdomeric regions is also observed in whole mount preparations of the adult eye in the mutants (C’) as compared to genetic controls (A’, B’). Rh1 staining is also more intense in *Df(3L) 218/+* (E-E’) as compared to its genetic control *cn, bw* (D-D’). (**G**, **F**) Increased Rh1 immunostaining intensity observed upon knockdown of *Prp31* by RNAi (G) as compared to its genetic background (F). Scale bars = 10µm. **(H)** Representative western blots for ß-Tubulin and Rhodopsin-1 from head lysates of *w*;;Prp31^P18^, st^1^/+* and its genetic background *w*\*;;st^1^/+. **(I)** Quantification from biological replicates (n=4) from western blotting. Bars represent the Rh1 levels calculated from intensity measurements of blots after normalization compared to that of loading control (Tubulin). On average, Rh1 levels are increased by 340% in *w*;;Prp31^P18^, st^1^/+* as compared to control, *w*;;st^1^/+*. This increase is evident despite the variability in the magnitude of increase.

All three deficiencies that remove the *Prp31* locus also exhibited increased Rh1 staining (Fig. 7E, E’ and Suppl. Fig. S5C, D) in comparison to the genetic control (Fig. 7D, D’). Finally, RNAi-mediated knockdown of *Prp31* also resulted in accumulation of Rh1 immunoreactivity (Fig. 7 F, G). Increased intensity of Rh1 immunostaining is due to increased levels of Rh1 as revealed by western blots of protein extracts isolated from adult heads (Fig. 7H, I). On average, Rh1 levels are increased by about four times in *Prp31*^*P18*^ heterozygous tissue as compared to tissue from genetic controls, *w*\*;;st^1^/+.

To determine whether rhodopsin accumulation contributes to light-dependent degeneration in *Prp31* mutant flies, we experimentally reduced rhodopsin levels by raising animals in carotenoid-free diet from embryonic stages onward. Carotenoids are precursors of the chromophore 11-cis-3-hydroxyretinal, which binds to opsin to generate the functional visual pigment rhodopsin in flies (Von LINTIG *et al.* 2010). In control genotypes, reduction of the chromophore halts maturation and ER to Golgi transport of rhodopsin, and an intermediate form accumulates in the perinuclear endoplasmic reticulum (Colley *et al.* 1991; Ozaki *et al.* 1993). Lack of dietary carotenoids strongly reduced Rh1 levels in the rhabdomere in controls and *Prp31* mutants and caused Rh1 accumulation in a peri-nuclear location (Fig. 8 A-D’).

**Figure 8:**
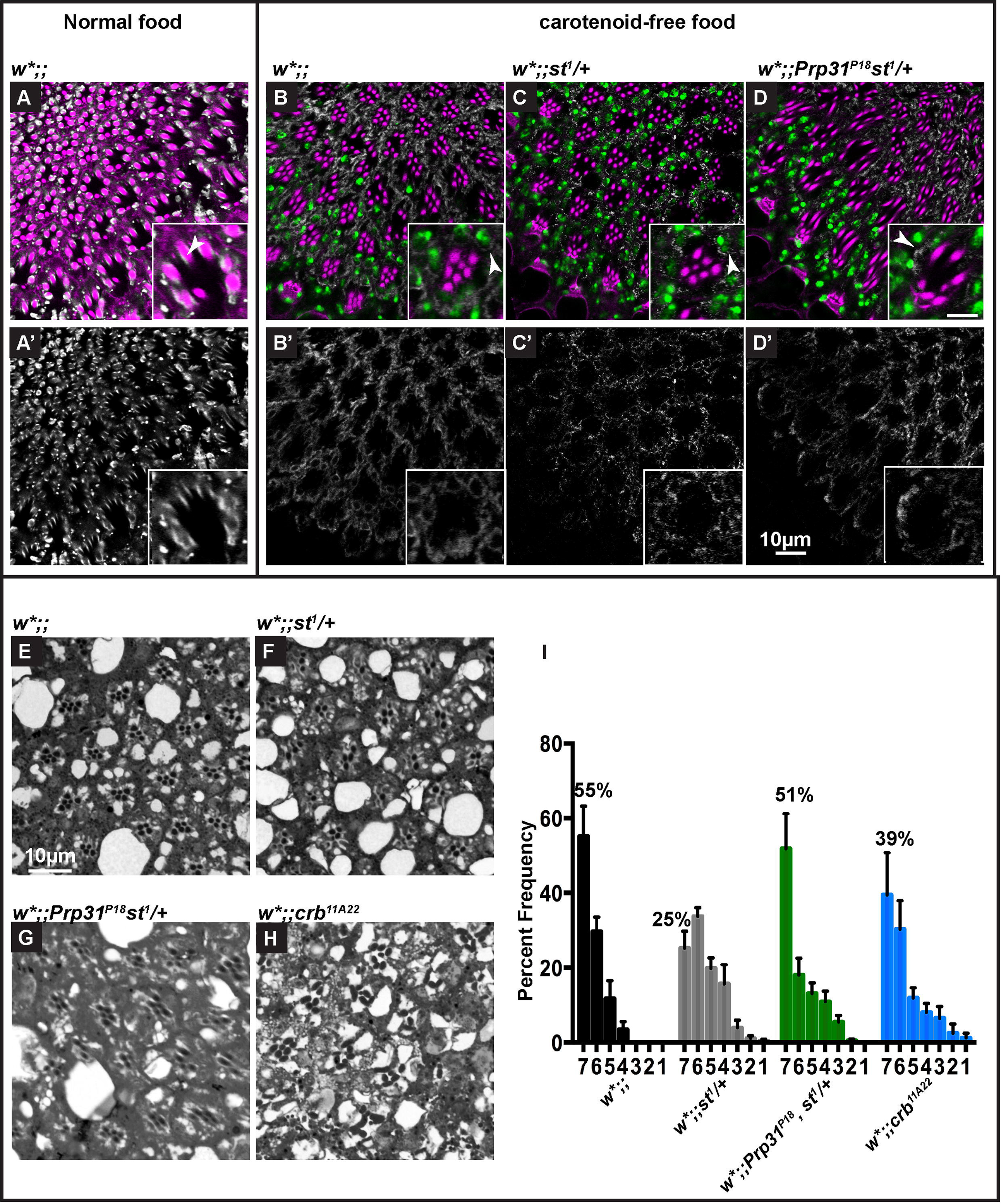
A carotenoid-depleted diet limits the extent of light-induced degeneration in hemizygous *Prp31* mutants. Representative images of 1µm confocal optical sections from 12µm cryosections of male eyes, of the genotypes indicated. Tissues are immunostained for Rh1 (white) and labelled with phalloidin (magenta) and DAPI (green), to stain the rhabdomeres and nuclei, respectively. (**A-D**) Overlay of all three channels, A’-D’ are images showing the extracted channel (Rh1). Insets show digital magnification of individual ommatidia. Reduction in Rh1 levels and change in its localization from the rhabdomeres to a peri-nuclear localization is observed when flies are fed a carotenoid-depleted diet (B-D’) as opposed to normal food (A-A’). Arrowheads indicate Rh1 localization in the rhabdomere (inset A-A’) as opposed to peri-nuclear localization (insets B-D’). Scale bar = 10µm. Inset scale bar = 5µm. (**E-H**) Representative bright-field images of Toluidine-blue stained semi-thin sections of eyes of *w\** (E), *w*;; st^1^/+*(F), *w*;;Prp31^P18^, st^1^/+* (G), and, *w*;;crb^11A22^* (H) adult males. Animals were raised on a carotenoid-depleted diet. Upon eclosion, they were aged for two days under regular light conditions and then subjected to a degeneration paradigm of exposure for 7 days to continuous, high-intensity light. Scale bar = 10µm. (**I**) Quantification of retinal degeneration as indicated by the number of surviving rhabdomeres observed upon high intensity, continuous light exposure. Bars represent mean ± s.e.m. (a minimum of n=60 ommatidia from eyes of 3 biological replicates) of the percent frequency of ommatidia displaying 1-7 rhabdomeres (Y-axis). Genotypes are indicated below.

Raising *Prp31* mutant animals in vitamin A depleted diet also suppressed light-dependent PRC degeneration. Under this dietary condition, more than 50% of ommatidia displayed 7 rhabdomeres, both in the control (*w*;;*) and in heterozygous *Prp31* flies (Fig. 8E-I). Interestingly, this dietary intervention did not suppress the degeneration observed in *w*;;st/+* eyes: only 25% ommatidia displayed the full complement of rhabdomeres. In agreement with previous reports (Satoh *et al.* 1998), the retinae of both genetic controls were more damaged when raised on carotenoid depleted diet as opposed to a standard diet (compare Fig. 2 and Fig. 8).

To conclude, these results point to Rh1 accumulation as a major cause of retinal degeneration in *Prp31* heterozygous flies.

### Mutations in *Prp31*^*P18*^ do not elicit increased oxidative stress signalling in photoreceptor cells

Although PRCs are specialised for light reception to initiate phototransduction, light at the same time is a stress factor and induces increased production of reactive oxygen species (ROS) (German *et al.* 2015). Increased levels of cellular ROS, in turn, induce antioxidant responses, which include expression of proteins against oxidative stress, e.g. superoxide dismutase (SOD) or glutathione S-transferase. Their activity can prevent cells from the detrimental consequences of oxidative stress, such as increased lipid oxidation or damage of proteins and DNA (Tomanek 2015). In photoreceptor cells, a failure of the antioxidant machinery to neutralise increased levels of ROS can lead to light-dependent retinal degeneration, for example in fly PRCs mutant for *crb* (Chartier *et al.* 2012).

This raised the question whether flies mutant for *Prp31* are subject to increased oxidative stress. Therefore, we analysed heterozygous *Prp31*^*P18*^ flies that carried the *gstD-GFP* reporter transgene. This reporter expresses GFP under the control of upstream regulatory sequences of *glutathione S-transferase* (*gstD1*), one of the genes involved in detoxification, whose expression is activated by oxidative stress (Sykiotis and Bohmann 2008). As shown previously, expression of this reporter correlates with the level of reactive oxygen species (ROS). This was revealed by comparing its activity with the signal induced upon application of a ROS-sensitive dye, Hydro-Cy3, in the midgut of adult flies stressed by feeding bacteria (Jones *et al.* 2013). Here, we examined GFP expression in-situ by immunostaining of adult mutant and control eye tissues, isolated form flies raised in regular light conditions. In control eyes (*gstD-GFP*/+), GFP expression was high in pigment and cone cells. Interestingly, barely any *gstD-GFP* expression was detected in PRCs (Fig. 9A). In eyes of *Prp31*^*P18*^/+ flies *gstD-GFP* expression was strongly increased in cone and pigment cells (Fig. 9B). Increased oxidative stress signalling in the retina of *Prp31*^*P18*^/+ flies was corroborated by using Dihydroethidium (DHE), a dye to detect ROS directly (Owusu-ANSAH *et al.* 2008) (Fig. 9D, E). Strikingly, the eyes of *Df(3L)217/+* flies (lacking one copy of the *Prp31* locus), did not show any increase in *gstD-GFP* expression nor in DHE staining (Fig. 9G-J), suggesting no altered ROS levels. Since *Prp31/+* flies are also heterozygous for *st*, we tested ROS levels in eyes of control flies with only one functional copy of *st*. Surprisingly, enhanced oxidative stress signalling and mild increase in ROS levels were observed in *w*;;st^1^/+* as compared to *w\** (Fig. 9C, F).

**Figure 9:**
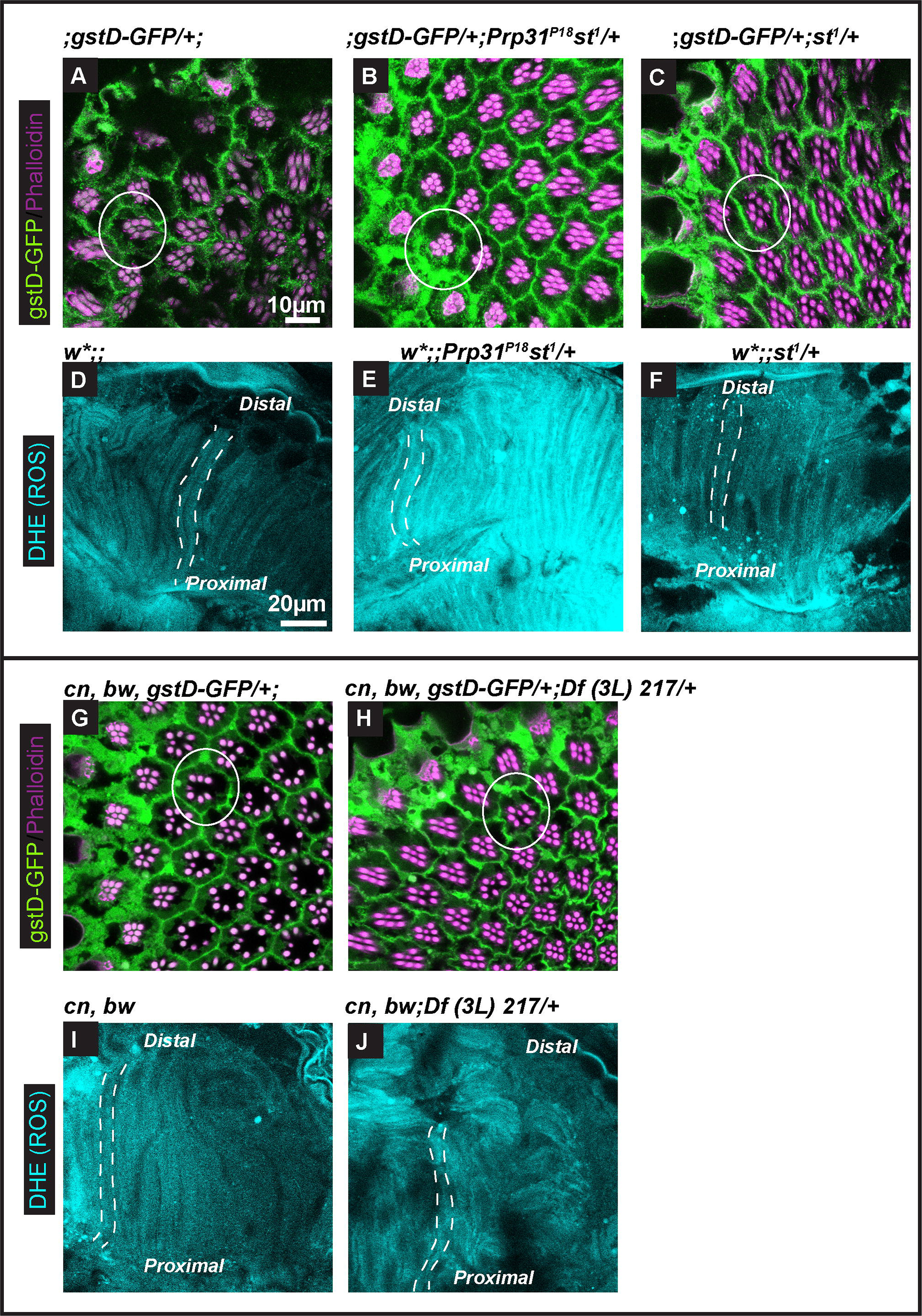
Increased oxidative stress signalling in eyes of *Prp31* heterozygous flies is due to *st* in the genetic background. (**A-J**) are images of 1µm confocal optical sections from 12µm cryosections (A-C, G-H) or whole tissue preparations (D-F, I-J) of eyes of adult flies raised in regular light conditions. Sections (A-C, G-H) are immunostained for anti-GFP (green) and phalloidin (magenta) for labelling *gstD* activity (oxidative stress signalling) and rhabdomeres, respectively. Whole tissue preparations (D-F, I-J) are labelled with Dihydroethidium (DHE), an indicator for the levels of reactive oxygen species (ROS). Individual ommatidia are outlined in white. Basal levels of oxidative stress signalling are observed in the pigment cells (surrounding ommatidia) in controls (A). This is enhanced in *Prp31*^*P18*^, *st*^1^/+(B) and *st*^1^/+ (C). Similarly, DHE levels are consistently increased in *w*;;Prp31*^*P18*^, *st*^1^/+ (E) and *w*;;st^1^/+* (F). No obvious increase in *gstD*-GFP staining and in DHE staining was observed in *Df/+* (H and J) as compared to the control (G and I).

From these results we conclude that loss of one copy of *Prp31* does not cause detectable increase in oxidative stress in PRCs, suggesting that increased accumulation of Rh1 in mutant PRCs is the major cause for retinal degeneration in this mutant.

## Discussion

Here we present a fly model for RP11, an autosomal-dominant human disease leading to blindness, caused by mutations in the splicing regulator PRPF31. Our results reveal that mutations in the *Drosophila* orthologue *Prp31* lead to PRC degeneration under light stress, thus mimicking features of RP11-associated symptoms. Similar as in human, mutations in *Drosophila Prp31* are haplo-insufficient and lead to retinal degeneration when hetero- or hemizygous. This is in stark contrast to mice heterozygous for *Prpf31*, which did not show any signs of PRC degeneration (Bujakowska *et al.* 2009), but rather late-onset defects in the retinal pigment epithelium (Graziotto *et al.* 2011; Farkas *et al.* 2014).

By using three different genetic approaches we provide convincing evidence that the knock-down of *Prp31* is the cause for the retinal degeneration observed. i) The two *Prp31* alleles induced by Tilling (*Prp31*^*P17*^ and *Prp31*^*P18*^) carry missense mutations in conserved amino acids of the coding region. ii) Flies heterozygous for any of three deletions, which remove the *Prp31* locus, exhibit the same phenotype. iii) RNAi-mediated knock-down of *Prp31* results in light-induced degeneration. From the results obtained we conclude that the two missense mutations mapped in *Prp31*^*P17*^ and *Prp31*^*P18*^ are strong hypomorphic alleles. First, the two *Drosophila* alleles characterized here are hemizygous and homozygous (in the case of *Prp31*^*P18*^) viable and fertile. Second, mutations in the two established *Prp31* fly lines are missense mutations, one located N-terminal to the NOSIC domain in *Prp31*^*P17*^ (G90R) and the other in the Nop domain in *Prp31*^*P18*^ (P277L) (see Fig. 1A), which most likely result in a reduced function of the respective protein. Whether protein levels are also decreased cannot be answered due to the lack of specific antibodies. In yeast, Prp31 is a component of the spliceosomal U4/U6 di-SNP, which contains, beside the base-paired U4 and U6 snRNAs, more than 10 other proteins, including Prp3 and Prp4. In this complex, Prp31 is required to stabilize a U4/U6 snRNA junction, which in turn is required for binding of Prp3/4 (Hardin *et al.* 2015). The Nop domain in human PRPF31 is involved in an essential step in the formation of the U4/U6-U5 tri-snRNP by building a complex of the U4 snRNA and a 15.5K protein. Consistent with this, many point mutations in human PRPF31, which are linked to RP11, have been mapped to the Nop domain. Mutations in amino acid H270 in the Nop domain of human PRPF31, for example, result in its reduced affinity to the complex formed by a stem-loop structure of the U4 snRNA and the 15.5K protein (Schultz *et al.* 2006; Liu *et al.* 2007). Interestingly, the mutated amino acid residue in *Drosophila Prp31^P18^* (P277L) lies next to a histidine (H278), which corresponds to amino acid H270 in the human protein (see Suppl. Fig. S1). Therefore, it is tempting to speculate that the *Drosophila* P277L mutation could similarly weaken, but not abolish the corresponding interaction of the mutant Prp31 protein with the U4/U6 complex. Further experiments are required to determine the functional consequences of the molecular lesions.

We noticed that the retinal phenotype observed upon reduction of *Prp31* is more variable than that observed upon loss of *crb* (see, for example, Fig. 2E) (Johnson *et al.* 2002; Spannl *et al.* 2017). This could be due to the fact that all *Prp31* conditions analyzed represent hypomorphic conditions with some residual function of the protein maintained. However, the expressivity of the mutant phenotype is not increased in *Prp31* hemizygous flies in comparison to that of *Prp31* heterozygous flies. This rather argues that the genetic background plays an important role. Background effects are often the result of the activity of so-called “modifier genes”, which modify the degree of the mutant phenotype due to their effects on the activity of the gene under discussion. This can be due either to a direct effect of the modifier on the functionality of the mutant allele (or the respective wild-type allele in a heterozygous condition), or to an indirect effect, e.g. as a result of a variation in a gene that acts in the same pathway as the gene under investigation. The availability of the so-called *Drosophila melanogaster* Genetic Reference Panel (DGRP) lines now allows to systematically screen for modifiers of a given mutation in about 200 inbred lines (Huang *et al.* 2014)[reviewed in (Mackay and Huang 2018)]. Using this library, modifiers of the locomotor defect in flies mutant for LRRK2 (leucine-rich kinase 2), a model for Parkinson’s disease, and for a Retinitis pigmentosa model based on defective rhodopsin (Chow *et al.* 2016; Lavoy *et al.* 2018), have been identified. Some of the candidates that affect the expressivity of the mutation studied are likely candidates to act in the same functional pathway as the respective disease gene. Interestingly, humans carrying the same molecular lesion in the *Prpf31* gene show an unusually high degree of phenotypic non-penetrance and can even be asymptomatic. Various causes have been uncovered to explain this feature, including a highly variable expression level of the wild-type *Prpf31* allele and changes in expression levels of trans-acting regulators (Rio FRIO *et al.* 2008) [reviewed in (Rose and Bhattacharya 2016)].

PRCs of flies lacking one functional copy of *Prp31* showed increased levels of Rh1 both in the rhabdomeres and in cytoplasmic punctae, as revealed by immunostaining and western blot analysis. Increased rhabdomeric Rh1, which, to our knowledge, has not been described for any other mutant, did not affect rhabdomere size or structure. This is different from observations in the mouse retina, in which transgenic overexpression of wild-type bovine or human rhodopsin induced an increase in outer segment volume of rod PRCs (Wen *et al.* 2009; Price *et al.* 2012). Increased Rh1 levels were also correlated to enhanced degeneration in *highroad* mutants. When analyzed in the presence of the folding-defective Rh1 allele, *ninaE^G69D^* to sensitize the background, PRC degeneration of *highroad* mutants was accelerated (Huang *et al.* 2018). Here, it has been hypothesized that *highroad,* encoding a carboxypeptidase, is required for Rh1 degradation. In several other *Drosophila* mutants, accumulation of Rh1 in endocytic compartments has been suggested to cause retinal degeneration due to its toxicity. For example, dominant mutations in *Drosophila ninaE* result in ER accumulation of misfolded Rh1 due to impaired protein maturation. This, in turn, causes an overproduction of ER cisternae and induces the unfolded protein response (UPR), which eventually leads to apoptosis of PRCs, both in flies and in mammals (Colley *et al.* 1995; Zhang *et al.* 2014; Kroeger *et al.* 2018). In the absence of carotenoids, rhodopsin maturation is impaired and opsin accumulates in perinuclear ER (Colley *et al.* 1991; Ozaki *et al.* 1993; Satoh *et al.* 1997).

Alternatively, as suggested for mutants in *norpA*, *arr2*, *rdgB* and *rdgC*, retinal degeneration can be induced by an abnormally stable, light-induced metarhodopsin-arrestin complex, which accumulates in the cytoplasm after endocytosis and is toxic (Alloway *et al.* 2000; Kiselev *et al.* 2000). Interestingly, mis-localisation of rhodopsin in human PRCs to sites other than the outer segment is a common characteristic of various forms of RP and is considered to contribute to the pathological severity (Hollingsworth and Gross 2012). Our data suggest that increased accumulation of rhodopsin causes degeneration in *Prp31* mutant retinas, since reduction of Rh1 by depletion of dietary carotenoid obliterated increased Rh1 immunoreactivity in *Prp31* mutant, caused opsin retention in perinuclear compartments and suppressed PRC degeneration. Currently, we cannot distinguish whether Rh1 accumulation in the rhabdomere or in the cytoplasm is responsible for light-dependent PRC degeneration. Our data further suggest that *Prp31* regulates, directly or indirectly, Rh1 levels at a posttranscriptional level, since no increase at the RNA level was detected in transcriptome analyses (own unpublished data). This is different from results obtained in primary retinal cell cultures, where expression of a mutant *Prpf31* gene reduced rhodopsin expression as a result of impaired splicing of the rhodopsin pre-mRNA (Yuan *et al.* 2005). It may be appealing to explore whether upregulation of Rh1 in *Drosophila Prp31* mutants is due to effects on the opsin protein, e. g. its stability, and/or the formation/stability of the chromophore. Additional defects could contribute to the mutant phenotype, such as impaired overall transcription or splicing defects, as described for *Prpf31* zebrafish models (Linder *et al.* 2011; Yin *et al.* 2011).

In several cases increased oxidative stress contributes to PRC degeneration, e. g. in PRCs mutant for *crb* (Chartier *et al.* 2012) or for *SdhA*, which encodes the succinate dehydrogenase flavoprotein subunit of mitochondrial complex II (Mast *et al.* 2008). Surprisingly, increase in ROS levels and ROS responses in the retina of w*; *Prp31*^*P18*^ st^1^ /+ flies could be traced back to the mutation in *st*, since the control *w*;;st^1^/+* showed higher levels of ROS as compared with *w\**, which correlates with enhanced retinal damage (Fig. 2, Suppl. Fig. S3). This defines *st^1^* as a dominant enhancer of *w\**, at least with respect to retinal degeneration. We would like to stress that all our analysis have been performed with *w\**, rather than with *w^1118^*, which is often used in comparable studies. Both alleles carry a big deletion, which includes the transcriptional and translational start site (Suppl. Fig. S1 and Suppl. Table 2). However, since retinal degeneration of *w^1118^* flies was much stronger under the light regime used here, all experiments and controls used the *w\** allele.

The enhancement of the *w* phenotype by *st^1^* seems surprising since both genotypes have unpigmented eyes. *w* and *st* encode members of the ATP binding cassette (ABC) transporters, and the White-Scarlet dimer is required for the transport of tryptophan, the precursor of xanthommatins (the brown pigments) into the granules of the eye’s pigment cells (Nolte 1950; Sullivan and Sullivan 1975; Tearle *et al.* 1989; Ewart and Howells 1998; Mackenzie *et al.* 1999; Mackenzie *et al.* 2000). In addition, *w* and *st* mutant flies have reduced numbers of capitate projections (Borycz *et al.* 2008), important specializations at the synapse of PRCs. Capitate projections are formed by finger-like invaginations of epithelial glia cells into the terminals of R1-R6 (Stark and Carlson 1986) and are sites of vesicle endocytosis and neurotransmitter recycling (Melzig *et al.* 1998; Fabian-Fine *et al.* 2003; Rahman *et al.* 2012). Reduced number of capitate projections were linked to retinal degeneration of *Drosophila* carrying mutations in *lin-7*, *cask* or *dlgS97*. Proteins encoded by these genes form a protein complex required in the postsynaptic lamina neurons to prevent retinal degeneration (Soukup *et al.* 2013). Whether *st^1^* also enhances the defects at the synapse of *w* remains to be elucidated. These results highlight the importance of carefully controlling the genetic background when studying retinal degeneration, including the choice of a specific allele.

## Acknowledgements

We would like to thank D. Bohmann for generously proving the *gstD-GFP* fly lines, the Bloomington Stock Centre for fly stocks, and the Developmental Studies Hybridoma Bank (DSHB) for antibodies. This work was supported by the fly facility, the light and electron microscopy facility and the sequencing facility of MPI-CBG. We thank K. Kapp (Univ. of Kassel, Germany) for technical advice on western blotting procedures, and K. Subramanian (MPI-CBG, Germany) for help with the spectrometer. This work was funded by the Max-Planck Society.

## References

Alloway, P. G., L. Howard and P. J. Dolph, 2000 The formation of stable rhodopsin-arrestin complexes induces apoptosis and photoreceptor cell degeneration. Neuron 28: 129–138.

Borycz, J., J. A. Borycz, A. Kubow, V. Lloyd and I. A. Meinertzhagen, 2008 Drosophila ABC transporter mutants white, brown and scarlet have altered contents and distribution of biogenic amines in the brain. J Exp Biol 211: 3454–3466.

Bujakowska, K., C. Maubaret, C. F. Chakarova, N. Tanimoto, S. C. Beck et al., 2009 Study of gene-targeted mouse models of splicing factor gene Prpf31 implicated in human autosomal dominant retinitis pigmentosa (RP). Invest Ophthalmol Vis Sci 50: 5927–5933.

Bulgakova, N. A., M. Rentsch and E. Knust, 2010 Antagonistic functions of two Stardust isoforms in Drosophila photoreceptor cells. Mol Biol Cell 21: 3915–3925.

Chartier, F. J.-M., E. J.-L. Hardy and P. Laprise, 2012 Crumbs limits oxidase-dependent signaling to maintain epithelial integrity and prevent photoreceptor cell death. J Cell Biol 198: 991–998.

Chen, X., H. Hall, J. P. Simpson, W. D. Leon-Salas, D. F. Ready et al., 2017 Cytochrome b5 protects photoreceptors from light stress-induced lipid peroxidation and retinal degeneration. NPJ Aging Mech Dis 3: 18.

Chinchore, Y., A. Mitra and P. J. Dolph, 2009 Accumulation of rhodopsin in late endosomes triggers photoreceptor cell degeneration. PLoS Genet 5: e1000377.

Chow, C. Y., K. J. Kelsey, M. F. Wolfner and A. G. Clark, 2016 Candidate genetic modifiers of retinitis pigmentosa identified by exploiting natural variation in Drosophila. Hum Mol Genet 25: 651–659.

Colley, N. J., E. K. Baker, M. A. Stamnes and C. S. Zuker, 1991 The cyclophilin homolog ninaA is required in the secretory pathway. Cell 67: 255–263.

Colley, N. J., J. A. Cassill, E. K. Baker and C. S. Zuker, 1995 Defective intracellular transport is the molecular basis of rhodopsin-dependent dominant retinal degeneration. Proc Natl Acad Sci U S A 92: 3070–3074.

Daiger, S. P., S. J. Bowne and L. S. Sullivan, 2014 Genes and Mutations Causing Autosomal Dominant Retinitis Pigmentosa. Cold Spring Harb Perspect Med 5.

Daiger, S. P., L. S. Sullivan and S. J. Bowne, 2013 Genes and mutations causing retinitis pigmentosa. Clin Genet 84: 132–141.

Dietzl, G., D. Chen, F. Schnorrer, K. C. Su, Y. Barinova et al., 2007 A genome-wide transgenic RNAi library for conditional gene inactivation in Drosophila. Nature 448: 151–156.

Ewart, G. D., and A. J. Howells, 1998 ABC transporters involved in transport of eye pigment precursors in Drosophila melanogaster. Methods Enzymol 292: 213–224.

Fabian-Fine, R., P. Verstreken, P. R. Hiesinger, J. A. Horne, R. Kostyleva et al., 2003 Endophilin promotes a late step in endocytosis at glial invaginations in Drosophila photoreceptor terminals. J Neurosci 23: 10732–10744.

Farkas, M. H., D. S. Lew, M. E. Sousa, K. Bujakowska, J. Chatagnon et al., 2014 Mutations in pre-mRNA processing factors 3, 8, and 31 cause dysfunction of the retinal pigment epithelium. Am J Pathol 184: 2641–2652.

Ferreiro, M. J., C. Perez, M. Marchesano, S. Ruiz, A. Caputi et al., 2018 Drosophila melanogaster White Mutant w(1118) Undergo Retinal Degeneration. Front Neurosci 11: 732.

German, O. L., D. L. Agnolazza, L. E. Politi and N. P. Rotstein, 2015 Light, lipids and photoreceptor survival: live or let die? Photochem Photobiol Sci 14: 1737–1753.

Graziotto, J. J., M. H. Farkas, K. Bujakowska, B. M. Deramaudt, Q. Zhang et al., 2011 Three gene-targeted mouse models of RNA splicing factor RP show late-onset RPE and retinal degeneration. Invest Ophthalmol Vis Sci 52: 190–198.

Hardin, J. W., C. Warnasooriya, Y. Kondo, K. Nagai and D. Rueda, 2015 Assembly and dynamics of the U4/U6 di-snRNP by single-molecule FRET. Nucleic Acids Res 43: 10963–10974.

Harris, W. A., W. S. Stark and J. A. Walker, 1976 Genetic dissection of the photoreceptor system in the compound eye of Drosophila melanogaster. J. Physiol. 256: 415–439.

Hollingsworth, T. J., and A. K. Gross, 2012 Defective trafficking of rhodopsin and its role in retinal degenerations. Int Rev Cell Mol Biol 293: 1–44.

Huang, H. W., B. Brown, J. Chung, P. M. Domingos and H. D. Ryoo, 2018 highroad Is a Carboxypetidase Induced by Retinoids to Clear Mutant Rhodopsin-1 in Drosophila Retinitis Pigmentosa Models. Cell Rep 22: 1384–1391.

Huang, W., A. Massouras, Y. Inoue, J. Peiffer, M. Ramia et al., 2014 Natural variation in genome architecture among 205 Drosophila melanogaster Genetic Reference Panel lines. Genome Res 24: 1193–1208.

Johnson, K., F. Grawe, N. Grzeschik and E. Knust, 2002 Drosophila Crumbs is required to inhibit light-induced photoreceptor degeneration. Curr Biol 12: 1675–1680.

Jones, R. M., L. Luo, C. S. Ardita, A. N. Richardson, Y. M. Kwon et al., 2013 Symbiotic lactobacilli stimulate gut epithelial proliferation via Nox-mediated generation of reactive oxygen species. EMBO J 32: 3017–3028.

Kiselev, A., M. Socolich, J. Vinós, R. W. Hardy, C. S. Zuker et al., 2000 A molecular pathway for light-dependent photoreceptor apoptosis in Drosophila. Neuron 28: 139–152.

Kroeger, H., W. C. Chiang, J. Felden, A. Nguyen and J. H. Lin, 2018 ER stress and unfolded protein response in ocular health and disease. FEBS J.

Kumar, J. P., and D. F. Ready, 1995 Rhodopsin plays an essential structural role in Drosophila photoreceptor development. Development 121: 4359–4370.

Lavoy, S., V. G. Chittoor-Vinod, C. Y. Chow and I. Martin, 2018 Genetic Modifiers of Neurodegeneration in a Drosophila Model of Parkinson’s Disease. Genetics.

Lee, Y. S., and R. W. Carthew, 2003 Making a better RNAi vector for Drosophila: use of intron spacers. Methods 30: 322–329.

Linder, B., H. Dill, A. Hirmer, J. Brocher, G. P. Lee et al., 2011 Systemic splicing factor deficiency causes tissue-specific defects: a zebrafish model for retinitis pigmentosa. Hum Mol Genet 20: 368–377.

Liu, M. M., and D. J. Zack, 2013 Alternative splicing and retinal degeneration. Clin Genet 84: 142–149.

Liu, S., P. Li, O. Dybkov, S. Nottrott, K. Hartmuth et al., 2007 Binding of the human Prp31 Nop domain to a composite RNA-protein platform in U4 snRNP. Science 316: 115–120.

Liu, W., Y. Xie, J. Ma, X. Luo, P. Nie et al., 2015 IBS: an illustrator for the presentation and visualization of biological sequences. Bioinformatics 31: 3359–3361.

Mackay, T. F. C., and W. Huang, 2018 Charting the genotype-phenotype map: lessons from the Drosophila melanogaster Genetic Reference Panel. Wiley Interdiscip Rev Dev Biol 7.

Mackenzie, S. M., M. R. Brooker, T. R. Gill, G. B. Cox, A. J. Howells et al., 1999 Mutations in the white gene of Drosophila melanogaster affecting ABC transporters that determine eye colouration. Biochim Biophys Acta 1419: 173–185.

Mackenzie, S. M., A. J. Howells, G. B. Cox and G. D. Ewart, 2000 Sub-cellular localisation of the white/scarlet ABC transporter to pigment granule membranes within the compound eye of Drosophila melanogaster. Genetica 108: 239–252.

Maita, H., H. Kitaura, T. J. Keen, C. F. Inglehearn, H. Ariga et al., 2004 PAP-1, the mutated gene underlying the RP9 form of dominant retinitis pigmentosa, is a splicing factor. Exp Cell Res 300: 283–296.

Mast, J. D., K. M. Tomalty, H. Vogel and T. R. Clandinin, 2008 Reactive oxygen species act remotely to cause synapse loss in a Drosophila model of developmental mitochondrial encephalopathy. Development 135: 2669–2679.

Melzig, J., M. Burg, M. Gruhn, W. L. Pak and E. Buchner, 1998 Selective histamine uptake rescues photo- and mechanoreceptor function of histidine decarboxylase-deficient Drosophila mutant. J Neurosci 18: 7160–7166.

Mishra, M., and E. Knust, 2013 Analysis of the Drosophila compound eye with light and electron microscopy. Methods Mol Biol 935: 161–182.

Mitra, A., Y. Chinchore, R. Kinser and P. J. Dolph, 2011 Characterization of two dominant alleles of the major rhodopsin-encoding gene ninaE in Drosophila. Mol Vis 17: 3224–3233.

Mordes, D., X. Luo, A. Kar, D. Kuo, L. Xu et al., 2006 Pre-mRNA splicing and retinitis pigmentosa. Mol Vis 12: 1259–1271.

Nolte, D. J., 1950 The eye-pigmentary system of Drosophila: the pigment cells. J Genet 50: 79–99.

Orem, N. R., X. L. and P. J. Dolph, 2006 An essential role for endocytosis of rhodopsin through interaction of visual arrestin with the AP-2 adaptor. J Cell Sci 119: 3141–3148.

Ostroy, S. E., M. Wilson and W. L. Pak, 1974 Drosophila rhodopsin: photochemistry, extraction and differences in the norp AP12 phototransduction mutant. Biochem Biophys Res Commun 59: 960–966.

Owusu-Ansah, E., A. Yavari, S. Mandal and U. Banerjee, 2008 Distinct mitochondrial retrograde signals control the G1-S cell cycle checkpoint. Nat Genet 40: 356–361.

Ozaki, K., H. Nagatani, M. Ozaki and F. Tokunaga, 1993 Maturation of major Drosophila rhodopsin, ninaE, requires chromophore 3-hydroxyretinal. Neuron 10: 1113–1119.

Parks, A. L., K. R. Cook, M. Belvin, N. A. Dompe, R. Fawcett et al., 2004 Systematic generation of high-resolution deletion coverage of the Drosophila melanogaster genome. Nat Genet 36: 288–292.

Pocha, S. M., A. Shevchenko and E. Knust, 2011 Crumbs regulates rhodopsin transport by interacting with and stabilizing myosin V. J Cell Biol 195: 827–838.

Poulos, M. G., R. Batra, K. Charizanis and M. S. Swanson, 2011 Developments in RNA splicing and disease. Cold Spring Harb Perspect Biol 3: a000778.

Price, B. A., I. M. Sandoval, F. Chan, R. Nichols, R. Roman-Sanchez et al., 2012 Rhodopsin gene expression determines rod outer segment size and rod cell resistance to a dominant-negative neurodegeneration mutant. PLoS One 7: e49889.

Rahman, M., H. Ham, X. Liu, Y. Sugiura, K. Orth et al., 2012 Visual neurotransmission in Drosophila requires expression of Fic in glial capitate projections. Nat Neurosci 15: 871–875.

Ray, P., X. Luo, E. J. Rao, A. Basha, E. A. Woodruff, 3rd et al., 2010 The splicing factor Prp31 is essential for photoreceptor development in Drosophila. Protein Cell 1: 267–274.

Rio Frio, T., N. Civic, A. Ransijn, J. S. Beckmann and C. Rivolta, 2008 Two trans-acting eQTLs modulate the penetrance of PRPF31 mutations. Hum Mol Genet 17: 3154–3165.

Rose, A. M., and S. S. Bhattacharya, 2016 Variant haploinsufficiency and phenotypic non-penetrance in PRPF31-associated retinitis pigmentosa. Clin Genet 90: 118–126.

Ruzickova, S., and D. Stanek, 2016 Mutations in spliceosomal proteins and retina degeneration. RNA Biol: 1–9.

Ryder, E., M. Ashburner, R. Bautista-Llacer, J. Drummond, J. Webster et al., 2007 The DrosDel deletion collection: a Drosophila genomewide chromosomal deficiency resource. Genetics 177: 615–629.

Satoh, A., F. Tokunaga, S. Kawamura and K. Ozaki, 1997 In situ inhibition of vesicle transport and protein processing in the dominant negative Rab1 mutant of Drosophila. J Cell Sci 110 (Pt 23): 2943–2953.

Satoh, A. K., H. Nagatani, F. Tokunaga, S. Kawamura and K. Ozaki, 1998 Rhodopsin Transport and Rab Expression in the Carotenpid-Deprived Drosophila melanogaster. Zoological Science 15: 651–659.

Satoh, A. K., and D. F. Ready, 2005 Arrestin1 mediates light-dependent rhodopsin endocytosis and cell survival. Curr Biol 15: 1722–1733.

Schindelin, J., I. Arganda-Carreras, E. Frise, V. Kaynig, M. Longair et al., 2012 Fiji: an open-source platform for biological-image analysis. Nat Methods 9: 676–682.

Schultz, A., S. Nottrott, K. Hartmuth and R. Luhrmann, 2006 RNA structural requirements for the association of the spliceosomal hPrp31 protein with the U4 and U4atac small nuclear ribonucleoproteins. J Biol Chem 281: 28278–28286.

Scotti, M. M., and M. S. Swanson, 2016 RNA mis-splicing in disease. Nat Rev Genet 17: 19–32.

Soukup, S. F., S. M. Pocha, M. Yuan and E. Knust, 2013 DLin-7 is required in postsynaptic lamina neurons to prevent light-induced photoreceptor degeneration in Drosophila. Curr Biol 23: 1349–1354.

Spannl, S., A. Kumichel, S. Hebbar, K. Kapp, M. Gonzalez-Gaitan et al., 2017 The Crumbs_C isoform of Drosophila shows tissue- and stage-specific expression and prevents light-dependent retinal degeneration. Biol Open 6: 165–175.

Stark, W. S., and S. D. Carlson, 1984 Blue and ultraviolet light induced damage to the Drosophila retina: ultrastructure. Curr Eye Res 3: 1441–1454.

Stark, W. S., and S. D. Carlson, 1986 Ultrastructure of capitate projections in the optic neuropil of Diptera. Cell Tissue Res 246: 481–486.

Stark, W. S., K. D. Walker and J. M. Eidel, 1985 Ultraviolet and blue light induced damage to the Drosophila retina: microspectrophotometry and electrophysiology. Curr Eye Res 4: 1059–1075.

Sullivan, D. T., and M. C. Sullivan, 1975 Transport defects as the physiological basis for eye color mutants of Drosophila melanogaster. Biochem Genet 13: 603–613.

Sun, Y., L. Liu, Y. Ben-Shahar, J. S. Jacobs, D. F. Eberl et al., 2009 TRPA channels distinguish gravity sensing from hearing in Johnston’s organ. Proc Natl Acad Sci U S A 106: 13606–13611.

Sykiotis, G. P., and D. Bohmann, 2008 Keap1/Nrf2 signaling regulates oxidative stress tolerance and lifespan in Drosophila. Dev Cell 14: 76–85.

Tanackovic, G., A. Ransijn, P. Thibault, S. Abou Elela, R. Klinck et al., 2011 PRPF mutations are associated with generalized defects in spliceosome formation and pre-mRNA splicing in patients with retinitis pigmentosa. Hum Mol Genet 20: 2116–2130.

Tearle, R. G., J. M. Belote, M. McKeown, B. S. Baker and A. J. Howells, 1989 Cloning and characterization of the scarlet gene of Drosophila melanogaster. Genetics 122: 595–606.

Tomanek, L., 2015 Proteomic responses to environmentally induced oxidative stress. J Exp Biol 218: 1867–1879.

von Lintig, J., P. D. Kiser, M. Golczak and K. Palczewski, 2010 The biochemical and structural basis for trans-to-cis isomerization of retinoids in the chemistry of vision. Trends Biochem Sci 35: 400–410.

Wang, S., K. L. Tan, M. A. Agosto, B. Xiong, S. Yamamoto et al., 2014 The retromer complex is required for rhodopsin recycling and its loss leads to photoreceptor degeneration. PLoS Biol 12: e1001847.

Wen, X. H., L. Shen, R. S. Brush, N. Michaud, M. R. Al-Ubaidi et al., 2009 Overexpression of rhodopsin alters the structure and photoresponse of rod photoreceptors. Biophys J 96: 939–950.

Will, C. L., and R. Luhrmann, 2011 Spliceosome structure and function. Cold Spring Harb Perspect Biol 3: pii: a003707.

Winkler, S., N. Gscheidel and M. Brand, 2011 Mutant generation in vertebrate model organisms by TILLING. Methods Mol Biol 770: 475–504.

Winkler, S., A. Schwabedissen, D. Backasch, C. Bokel, C. Seidel et al., 2005 Target-selected mutant screen by TILLING in Drosophila. Genome Res 15: 718–723.

Xiong, B., V. Bayat, M. Jaiswal, K. Zhang, H. Sandoval et al., 2012 Crag is a GEF for Rab11 required for rhodopsin trafficking and maintenance of adult photoreceptor cells. PLoS Biol 10: e1001438.

Xiong, B., and H. J. Bellen, 2013 Rhodopsin homeostasis and retinal degeneration: lessons from the fly. Trends Neurosci 36: 652–660.

Yin, J., J. Brocher, U. Fischer and C. Winkler, 2011 Mutant Prpf31 causes pre-mRNA splicing defects and rod photoreceptor cell degeneration in a zebrafish model for Retinitis pigmentosa. Mol Neurodegener 6: 56.

Yuan, L., M. Kawada, N. Havlioglu, H. Tang and J. Y. Wu, 2005 Mutations in PRPF31 inhibit pre-mRNA splicing of rhodopsin gene and cause apoptosis of retinal cells. J Neurosci 25: 748–757.

Zhang, S. X., E. Sanders, S. J. Fliesler and J. J. Wang, 2014 Endoplasmic reticulum stress and the unfolded protein responses in retinal degeneration. Exp Eye Res 125: 30–40.

Zhao, C., D. L. Bellur, S. Lu, F. Zhao, M. A. Grassi et al., 2009 Autosomal-dominant retinitis pigmentosa caused by a mutation in SNRNP200, a gene required for unwinding of U4/U6 snRNAs. Am J Hum Genet 85: 617–627.

